# Chromatin accessibility variations across pancreatic islet maturation

**DOI:** 10.1101/782318

**Authors:** Jonathan Sobel, Claudiane Guay, Adriana Rodriguez-Trejo, Lisa Stoll, Véronique Menoud, Romano Regazzi

## Abstract

Glucose-induced insulin secretion, a peculiar property of fully mature *β*-cells, is only achieved after birth and is preceded by a phase of intense proliferation. These events occurring in the neonatal period are decisive for the establishment of an appropriate functional *β*-cell mass that provides the required insulin throughout life. However, key regulators of gene expression involved in cellular reprogramming along pancreatic islet maturation remain to be elucidated. The present study addressed this issue by mapping open chromatin regions in newborn versus adult rat islets using the ATAC-seq assay. Accessible regions were then correlated with the expression profiles of mRNAs to unveil the regulatory networks governing functional islet maturation. This led to the identification of Scrt1, a novel transcriptional repressor controlling *β*-cell proliferation.

## Introduction

Pancreatic *β*-cells are highly specialized cells displaying the unique functional feature to release insulin in response to glucose and other stimuli. Incapacity of *β*-cells to secrete the appropriate amount of insulin to cover the organism’s needs leads to diabetes mellitus development. *β*-cells acquire the ability to secrete insulin through a poorly understood postnatal maturation process implying transcriptional reprogramming of gene and non-coding RNA expression that terminates only after weaning (1, 2). Cells that generate and secrete insulin can be produced *in vitro* using various methods (3, 4) and are similar to *β*-cells in many aspects. However, these cells show lower glucose-stimulated insulin secretion (GSIS) and transcriptome differences compared with normal *β*-cells (5). Consequently, there is a need to better understand the regulation of the maturation process to enable the engineering of fully functional surrogate insulin-producing cells. Likewise, the transcriptional control of pancreas development and islet differentiation is still subject of intensive investigations (6–8). In the past decade, the emergence of various next-generation sequencing technologies and experimental procedures allowed querying epigenome and gene expression profiles with an unprecedented precision at a genome-wide or transcriptome-wide scale. Indeed, a recent study by Ackermann *et al.* identified human *α* and *β*-cell specific mRNAs and regulatory elements using RNA-seq and ATAC-seq, respectively (9).The ATAC-seq method probes DNA accessibility with a hyperactive Tn5 transposase, which inserts sequencing adapters into accessible chromatin regions. Sequencing reads can then be used to infer regions of increased accessibility, as well as to map regions of transcription factor binding and nucleosome position (10). This allows the identification of transcription factors (TFs) that will bind to specific binding sites (TFBS) located in an open chromatin region, which can be close or distant to the transcription start site (proximal/distal regulation). Moreover, sets of TFs can cooperate on cis-regulatory modules (CRM), which are DNA stretches of about 100-1000 bp, to produce specific regulatory events (11). Taking advantage of this approach, we aimed at determining how neonatal islet maturation is controlled at the transcriptional level and to identify the key transcription factors involved. We thus used ATAC-seq to search for open chromatin regions in the islets of 10-day-old (P10) and 3-month-old (adult) rats. Using this strategy, we found about 100’000 putative regulatory regions, 20% of which displayed significant accessibility changes upon maturation. We then used two different computational approaches to investigate potential TFBS motifs in the regions with accessibility changes. This allowed us to identify putative regulatory elements in the promoter or in distal CRM nearby differentially expressed mRNAs that may be implicated in islet maturation. As a result, we obtained a global picture of the transcriptional events taking place during pancreatic islet maturation. Moreover, we experimentally confirmed the involvement of a novel transcriptional regulator, Scrt1, in the neonatal maturation of pancreatic *β*-cells.

## Results

To study the transcriptional regulation of pancreatic islet maturation, we performed high throughput sequencing of Transposase-Accessible Chromatin of 10-day-old and adult rat islets (Figure 1A). We then applied bioinformatic methods for quality control, peak detection, differential accessibility analysis and motif finding (Figure 1B). For example, the Magnesium Transporter 2 (*Mrs2*) locus depicts two prominent ATAC-seq signal peaks, and the 3’ end peak shows an important increase of accessibility after the maturation process. Quality control of the samples (table 1) showed an average of 370 million reads sequenced with 94.3% of reads with a MAPQ score above 30. The fragment size analysis showed the expected profile of ATAC-seq (Supplementary figure S1A) with usual oscillations due to the presence of nucleosomes (∼150 bp) or side accessibility (∼10 bp) (12). In addition, adult and P10 samples were well clustered, when evaluated by the correlation between samples (Supplementary figure S1B) or using the first PCA component (Supplementary figure S1C). To detect accessible sites (ACS), we performed a peak calling using MACS2 (13) (Methods), leading to the detection of ∼ 102000 ACS (Supporting Information (1)). These sites were quantified in each sample separately and annotated with the closest transcription start site (TSS). Moreover, a differential accessibility analysis was performed using EdgeR (15) (Figure 2), and the values of the analysis reported in the source data (1). ACS were divided in 3 groups: Stable, significantly more accessible in P10 (Down) and significantly more accessible in adults (Up), with p-value < 0.05 and FDR < 0.2 (Figure 2A). About 20% of the ACS showed differential accessibility upon maturation with 11.8% of down ACS and 7.1% of up ACS (Figure 2B,C).

**Table 1.**
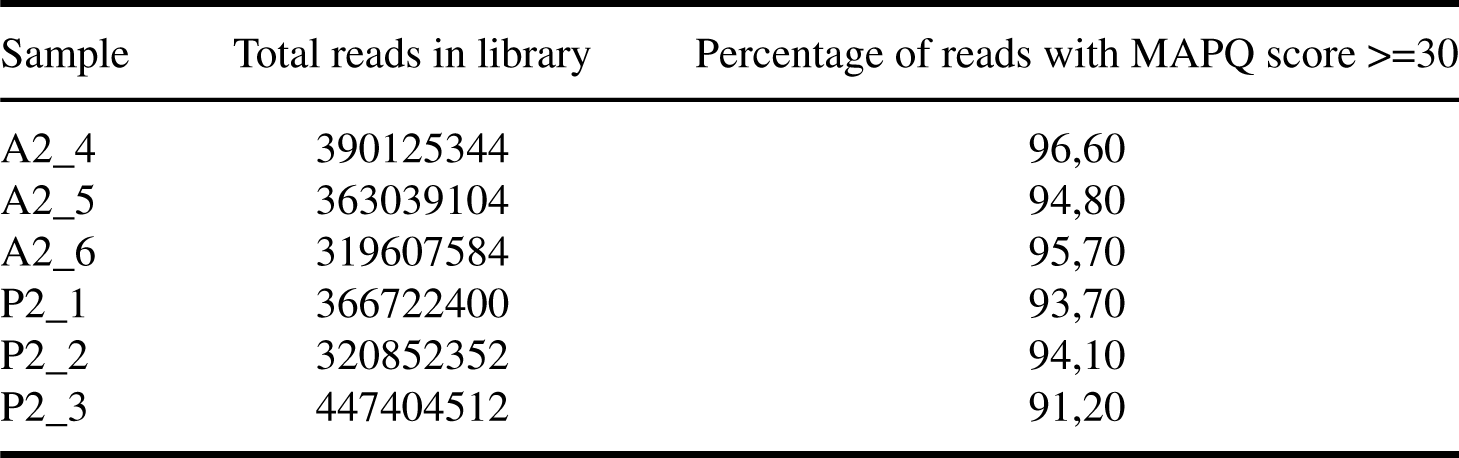
ATAC-seq libraries quality control

**Fig. 1.**
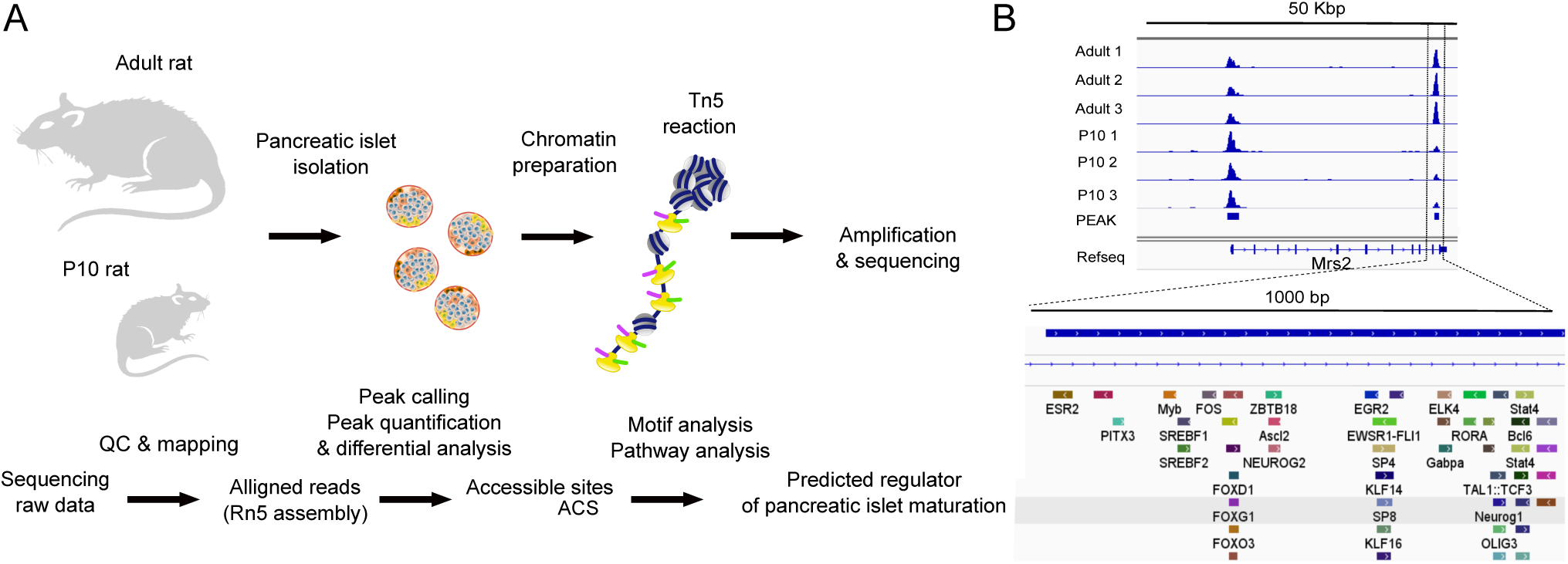
ATAC-seq successfully identified accessible sites (ACS) associated with pancreatic islet maturation. (A) Summary of the ATAC-seq experimental design and computational analysis pipeline. We extracted nuclei from the isolated islets of 3 adult (3 m.o.) rats and 3 litters of rat pups at postnatal day 10 (P10) to perform the Tn5 reaction as described in (12) and prepare the library for sequencing. The computational pipeline involved a quality control of the sequencing data followed by read alignment to the rat reference genome (Rn5 assembly). ACS were identified using the peak calling tool MACS2 (13) and quantified for each sample separately. The ACS sequences were scanned using FIMO (14) and analyzed to identify TFBS motifs that are implicated in the islet maturation process and in related pathways (See methods). (B) Example of identified ACS. The ACS nearby *Mrs2* transcription end site is significantly higher in adult rats. This ACS contains several motifs of TFBS. See also Figure S1 and table 1

**Fig. 2.**
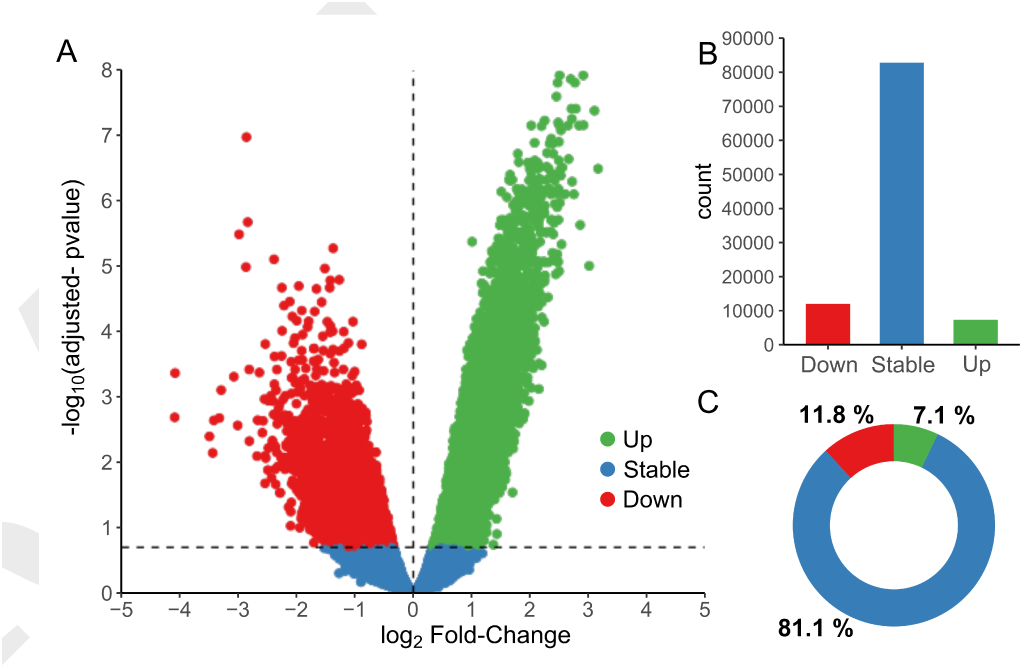
Differential analysis of Trn5 integrations in accessible sites revealed that ∼20% of the ACS were significantly changing during islet maturation. (A) Volcano plot representation of the log_2_ fold-change of Trn5 integration between 10-day-old pups (P10) and adult rat islets in the x-axis, and the FDR adjusted p-value in the y-axis. ACS significantly more accessible in adults are represented in green (Up), those more accessible in P10 in red (Down), and those remaining stable in blue (p-value <0.05, FDR <0.2, n=3). ACS count in (B) and percentage in (C) for each group (Up, Stable and Down). See also source data (1) and supplementary figures S2,S3 and S4

### Promoter and distal ACS depicted distinct accessibility patterns along islet maturation

To assess the impact of the ACS location in respect to the closest gene, we used ChIPseeker (16) to annotate our ACS and 100’000 randomly selected sites (Supplementary figure S2A). We observed that true ACS were enriched for exonic and intronic sequences and transcript start sites (Supplementary figure S2B). Interestingly, a larger fraction of sites more accessible in adults (Up) belonged to the distal class and were located on introns while ACS with decreased accessibility in adults (Down) were enriched in promoters. The localization analysis showed that 2/3 of the ACSs were distal sites and about 10% were located in the TSS or proximal class. We then looked at all the transcription start sites (TSS) and the transcription end sites (TES) of genes annotated with one or more significantly changing ACS (Supplementary figure S3). We observed that TSS of genes nearby ACS more accessible in adults showed a precise start signal, while the other group seemed to have larger accessible sites, maybe due to less constrained transcription (wide promoters or bi-directional promoters (17)). In addition, we observed that ACS less accessible in adults showed a drastic decrease of the signal, indicating that these sites are closing due to chromatin remodelling events. On the other hand, in ACS that showed a higher accessibility in adults, the difference between P10 and adult sites was not as substantial, suggesting that these ACS were still open in P10 but potentially bound by a different set of transcription factors.

### ACS significantly affected were located nearby genes involved in islet maturation

Next, we investigated if ACS changing in P10 versus adult rat islets control the expression of genes displaying significant differences upon maturation (2). Of the 19311 ACS differentially accessible with p-value below 0.05 and FDR below 0.2, 11372 (∼60%) were located in the vicinity of differentially expressed genes. These ACS were annotated as enhancers or repressors, depending whether their accessibility was respectively correlated or anti-correlated with the expression changes in the nearby gene (Supplementary figure S4). We found that ACS less accessible in adult (Down) were enriched nearby TSS (down-regulated gene enhancer and up-regulated gene repressor). In addition, several KEGG pathways were enriched in adult samples such as insulin secretion, circadian rhythm, and calcium signaling, while carbon metabolism, PI3-Akt, and proliferation related annotations (cancer) were enriched in P10 samples. In most cases, both enhancer and repressor ACS contribute to the output gene expression. Thus, a gene might be regulated by several ACS, some bound by enhancer proteins, and others targeted by repressors. Consequently, the ATAC-seq signal of specific ACS may be increased, while the nearby gene is repressed.

### ACS display enhancer activity

Next, we experimentally tested if the identified ACS nearby genes important for proper pancreatic *β*-cell function (18) display enhancer activity. For this purpose we cloned 5 ACS sequences (Supplementary Table (2), Figure 3) near *Syt4, Pax6* (two ACS), *Mafb* and *NeuroD1* in a luciferase reporter construct driven by a minimal promoter. Interestingly, the inclusion of the ACS close to *Syt4, MafB*, and *NeuroD1* resulted in an increase in luciferase activity, compared to the empty pGL3 vector. In addition, mRNA expression of *NeuroD1* and *MafB*, tested by qPCR, confirmed the higher expression of these genes in 10-day-old pups. Thus, we concluded that the identified ACS are likely to be involved in transcriptional regulation of nearby genes.

**Fig. 3.**
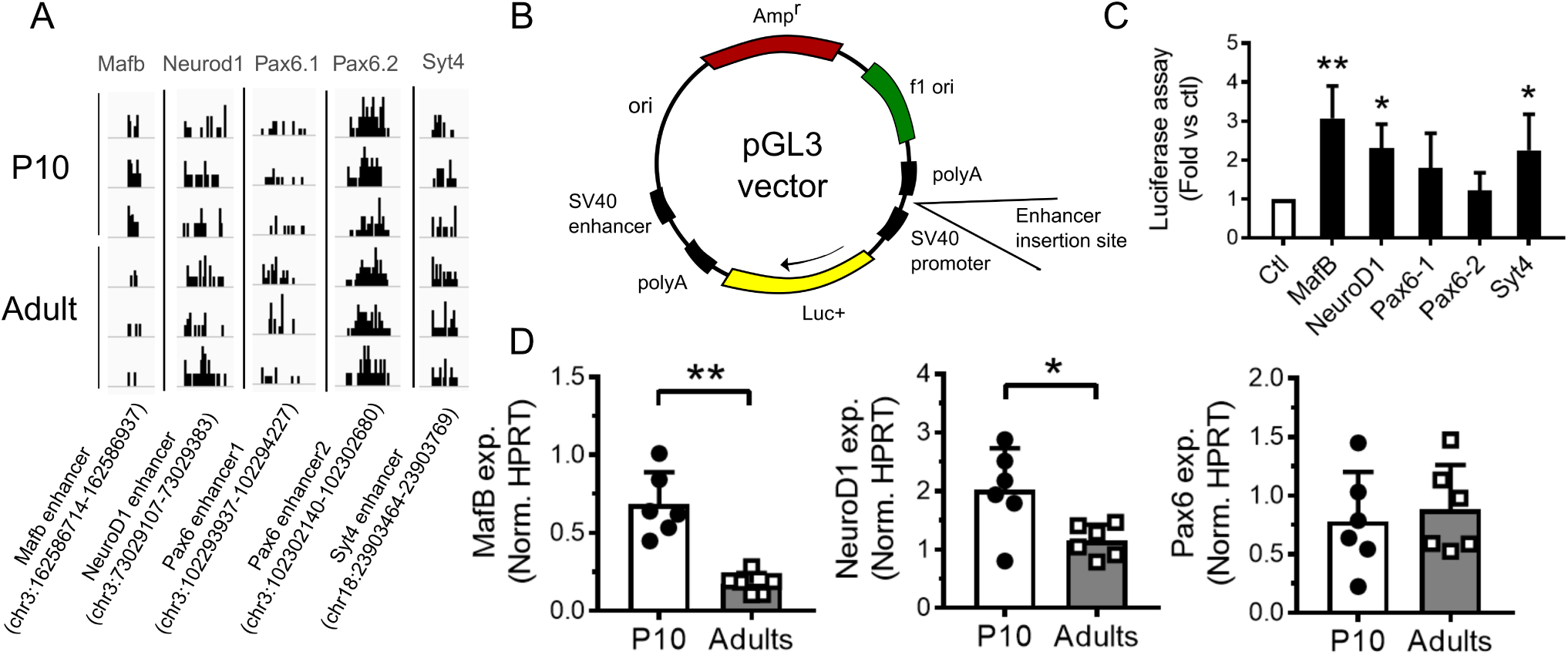
Enhancer activity assessment using a luciferase assay. (A) Trn5 integrations on *Mafb, Syt4, Neurod1* and two *Pax6* accessible sites. (B) Scheme of pGL3 vector used for the luciferase assay. (C) Luciferase activity was measured in INS 832/13 cells transfected with a pGL3 empty vector (Ctl) or a pGL3 vector containing an enhancer region for the indicated gene (*Mafb, NeuroD1, Pax6* and *Syt4*). Results are expressed as fold change versus control. (D) Gene expression in P10 and adult rat islets were measured by qPCR and normalized to the housekeeping gene *Hprt1. Syt4* gene expression is available in the figure 6. * p < 0.05, * * p < 0.01 by Student’s t-test or by one-way Anova, Dunnett’s post-hoc test. See also Source data (2).

### Identification of transcriptional regulators and chromatin remodelers of pancreatic islet maturation

Accessibility analysis enables the detection of transcription factors (TFs) or DNA binding motifs (TFBS) affecting the chromatin state and the transcription of nearby genes. In order to decipher the combinatorial code of transcription factor binding sites that allow islet maturation, we scanned the sequence of each ACS using FIMO (14) together with the position weight matrices of Jaspar 2016 (19). With these sequence scans we could perform a motif set enrichment analysis for the accessibility in P10 or in adult islet cells using the FGSEA algorithm (20) (Supplementary Table (3), Figure 4A,B). Thus, using the set of significantly changing ACS and their respective accessibility log_2_ Fold Change, we were able to identify TFBS motifs that were either enriched in P10 or in adult rat islets. We found several TFBS that were previously implicated in islet maturation such as MAF, FOX, FOS/JUN, NRF, and E2F (6). Moreover, we observed the enrichment of several motifs recognized by transcriptional repressors and insulators such as SCRT1 or CTCF. To confirm these results and detect additional motifs playing a role in islet maturation, we used a more sensitive method. We applied a penalized linear model GLMnet (21) to all ACS with the matrix of motif match as predictors and the log_2_ fold-change as output vector. With this method, we could identify a set of TFBS motifs susceptible to play an important role in the maturation and we could compute an activity (*β*) for each motif (Supplementary Table (4),Supplementary Table S5). This permitted to confirm most of the hits discovered using the FGSEA and to detect other TFBS motifs such as RFX, SREB, NKX6, REL, MEIS, and TEAD3.

**Fig. 4.**
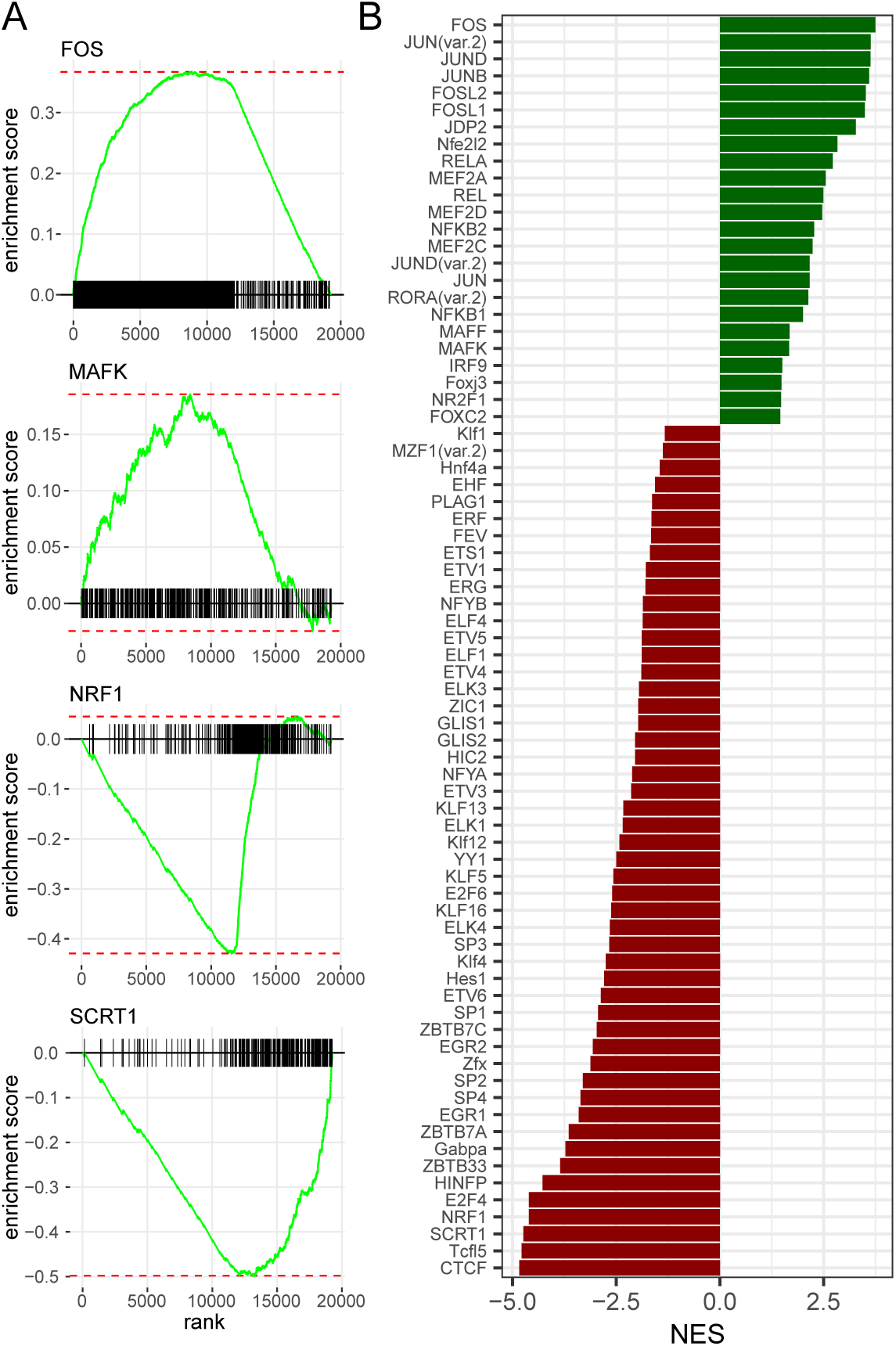
Motif enrichment analysis in ACS identified putative regulators of postnatal pancreatic islet maturation.(A) We scanned every ACS sequence using FIMO (14) and the Jaspar core vertebrate (19) position-weight matrices (PWMs) to construct the motif-ACS annotation. We used this annotation and the log_2_ fold change of significantly modified ACS with the FGSEA algorithm (20) to find putative regulators of the maturation process. The FGSEA algorithm calculates an enrichment score for each binding site motif by evaluating its distribution within the list of ACS ordered by accessibility log_2_ fold-change. Figure represents four enriched TFBS motifs (B) FGSEA Normalized Enrichment Score for top enriched motifs (FGSEA adjusted p-value < 0.05). See also supplementary figure S5 and supplementary data set (3) and (4).

### Scrt1 represses *β*-cell proliferation

We next focused on Scrt1, a transcriptional repressor involved in neuroendocrine development (22, 23) but whose function has not yet been investigated in *β*-cells. Binding sites for this transcriptional repressor were highly enriched in the chromatin regions that close upon *β*-cell maturation. In agreement with the lower chromatin accessibility for Scrt1 binding sites, *Scrt1* expression is increased in adult islets (Figure 5A). Transfection of adult rat islet cells with a set of siRNAs targeting *Scrt1* led to a decrease in the expression of the repressor of about 70%. (Figure 5B). Down-regulation of *Scrt1* did neither affect insulin secretion in response to glucose (Figure 5C) nor insulin content (Figure 5D). Apoptosis measured by Tunel assay in both control condition or in response to pro-inflammatory cytokines was also not affected (Figure 5E). Interestingly, knockdown of *Scrt1* in adult *β*-cells to levels similar to those present in newborn rats resulted in a rise in proliferation, suggesting an involvement of this transcriptional repressor in postnatal *β*-cell mass expansion (Figure 5F).

**Fig. 5.**
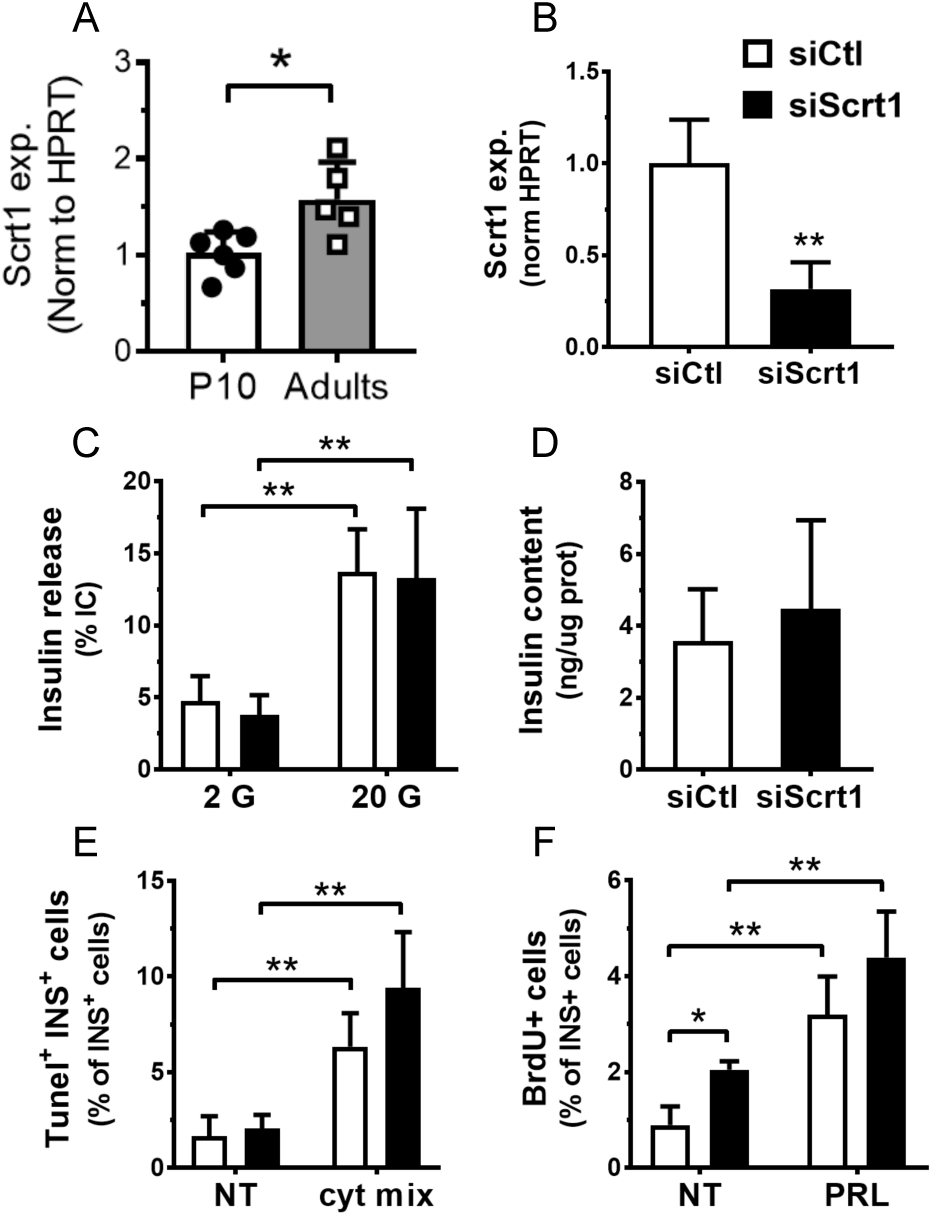
Downregulation of *Scrt1* promotes *β*-cell proliferation. A) *Scrt1* expression is increased in adult versus P10 rat islets. B-F) Dispersed adult rat islet cells were transfected with a control siRNA (siCtl) or siRNAs directed against *Scrt1* (siScrt1). Experiments were performed 48h post-transfection. B) siScrt1 led to a 70% decrease in *Scrt1* expression as measured by qPCR and normalized to *Hprt1* house-keeping gene levels. C) Insulin release in response to 2 or 20 mM glucose (G) and D) insulin content were determined by ELISA. E) Apoptosis of insulin positive cells was assessed using Tunel assay in basal (NT) condition or in response to a mix (cyt mix) of pro-inflammatory cytokines (IL-1*β*, TNF-*α*, IFN-*γ*). F) The fraction of proliferative insulin positive cells was determined by BrdU incorporation in basal (NT) or stimulated (prolactin, PRL) conditions. * p < 0.05, * * p < 0.01 by Student’s t-test or by one-way Anova, Tukey’s post-hoc test.

### Identification of Scrt1 targets involved in maturation

As Scrt1 is regulating the proliferative capacity of *β*-cells, we next aimed at finding its targets. For this purpose, we FACS-sorted adult rat islet cells to separate *α*-cells and *β*-cells (Supplementary FigureS6). We observed that *α*-cells and *β*-cells express *Scrt1* at similar levels. Subsequently, we performed RNA-seq on *β*-cells exposed to a control siRNA or to siScrt1 (Figure 6). Differential expression analysis between siScrt1 and control samples revealed that 168 genes were significantly impacted by silencing *Scrt1* with a FDR adjusted p-value below 0.05 (Figure 6A). Of these 168 genes, 111 were down-regulated and 57 were up-regulated. As expected, among the potential targets of Scrt1 we found genes related to proliferation such as *Notch1, Parp16, Ppp3r1, Ppp2r1b* and *Ywhag* (Supplementary figure S7 A,B). Moreover, some genes related to glucose signaling and GSIS such as *Syt4*, or to sphingolipid metabolism as *Ugt8* were down-regulated when knocking down *Scrt1*. Then, we compared the set of genes affected by *Scrt1* silencing and the ones differentially expressed upon maturation (in postnatal 10-day-old (P10) versus adult rat islets (2, 24)). We found a common set of 62 genes changing in both data sets with a FDR adjusted p-value below 0.05 (Figure 6B, Supplementary table (7)). Interestingly, we observed a significant anti-correlation (correlation test p-value= 0.013) between the fold-changes from the comparison between siScrt1 versus siCtl in adult rat *β*-cells and P10 versus adult islets. We confirmed using qPCR that *NFATc1, NFATc2, Notch1*, and *Syt4* were controlled by Scrt1 and were inversely changing upon maturation (Figure 6C,D). In addition, a gene ontology enrichment analysis for biological processes revealed that autophagy and oxygen sensing are over represented in these 62 genes (Supplementary table (8)). Overall, our results suggest that *Scrt1* is implicated in the switch between the proliferative state and the fully functional state of *β*-cells along pancreatic islet maturation.

**Fig. 6.**
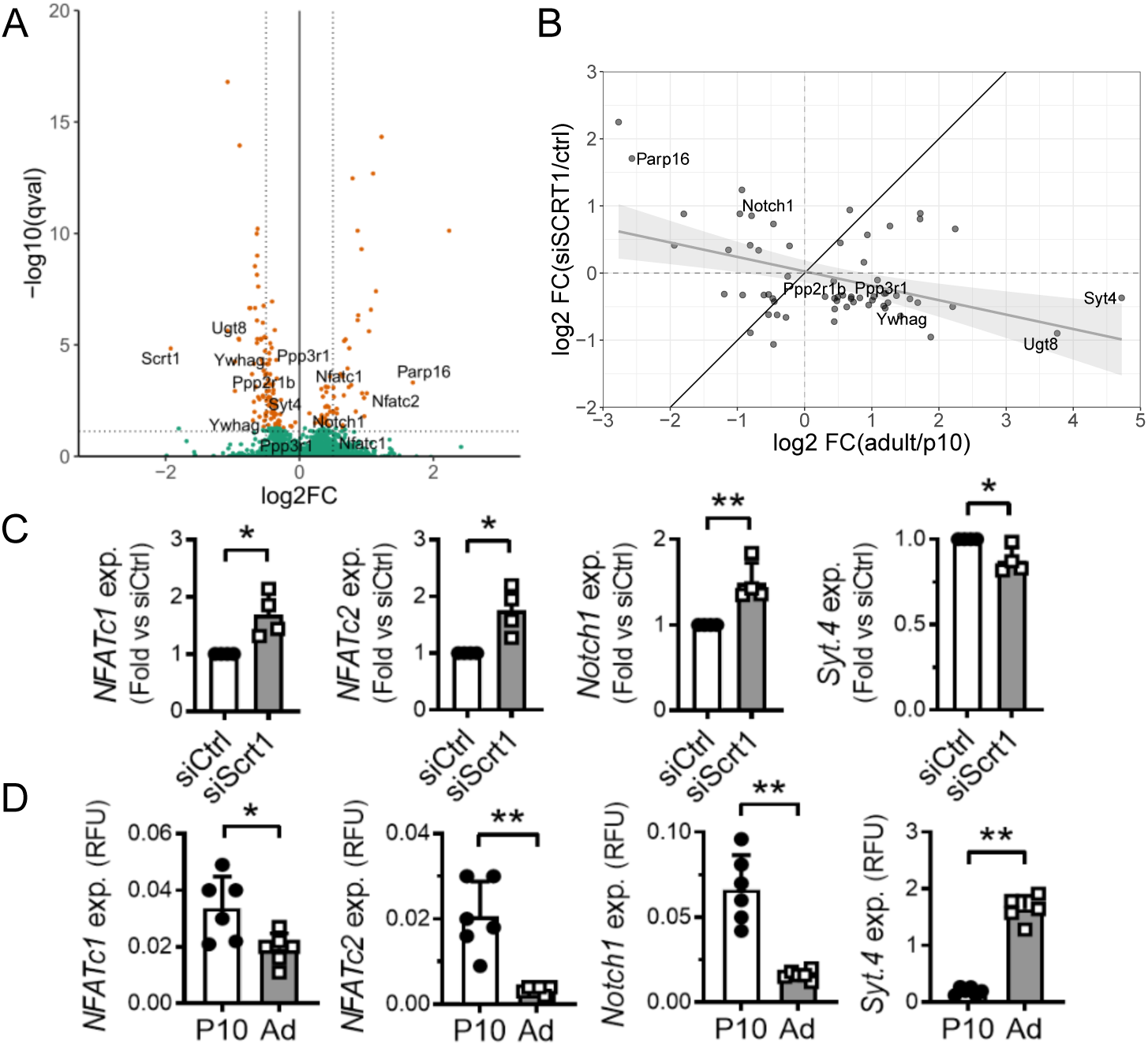
Changes induced by silencing Scrt1 were anti-correlated with the maturation signature of Scrt1 targets. FACS-sorted adult rat *β*-cells were transfected with a control siRNA (siCtl) or siRNAs directed against *Scrt1* (siScrt1).RNA-extraction and library preparation for RNA-seq were performed 48h post-transfection. A) Volcano plot of gene expression changes induced by *Scrt1* knockdown. B) Scatter plot of log_2_ fold changes of differentially expressed genes from adult/P10 measured by micro-array (n=3) versus log_2_ fold changes of siScrt1/control measured by RNA-seq (n=5). C) qPCR confirmation of gene expression in FACS-sorted *β*-cells transfected with siCtrl or siScrt1, D) P10 versus adult rat islets. Student T test, * p< 0.05, * * p< 0.01. see also supplementary Figure S6, S7 and Supplementary Tables (5),(6),(7),(8).

## Discussion

Postnatal islet maturation is a critical process to achieve proper *β*-cell function. Immediately after birth, *β*-cells are not fully functional and have to undergo a major gene reprogramming to acquire the ability to secrete adequate amounts of insulin in response to glucose (1). Our group and others (2, 5, 24) have shown a large-scale rewiring of transcriptional programs occurring during the neonatal period. However, little is known on cis-regulation of gene expression at the chromatin level before and after weaning. Chromatin accessibility of human islets on a genome-wide scale has been previously produced using FAIRE-seq (18, 25). More recently, several studies took advantage of ATAC-seq together with GWAS to identify causal variants of T2D in cis-regulatory elements in human (26–29). Another study identified cell-type specific accessible sites and transcription factor binding sites in *α, β* and acinar cells (9). However, the transcriptional regulation of postnatal islet maturation at the chromatin level has not been reported so far. In this project, we employed ATAC-seq (12) to produce a global map of accessible sites (ACS) in the islets of 10-day-old pups and in adult rats. This permitted to detected more than 100’000 ACS, among which about 20% were differentially accessible before and after the functional maturation of *β*-cells. Interestingly, one of the ACS with the most significant p-value and larger fold-change was located on the 3’UTR of the *Mrs2* gene. This gene is encoding a magnesium transporter at the surface of the mitochondria (30). Genetic variants in this magnesium-related ion channel were previously associated with type 2 diabetes and pancreatic cancer (31, 32). Next, we integrated these accessibility maps with mRNA micro-array data from (2, 24) in the same model and we found that ∼ 60% of the accessible sites were nearby differentially expressed genes, suggesting that these cis-regulatory elements are involved in the transcriptional regulation of gene expression. Several pathways previously related to the maturation process such as insulin secretion or circadian rhythms are under the control of these accessible cis-regulatory elements. Interestingly, several components of the core clock are not rhythmic in 10 days old pups but are consistently oscillating in *β*-cells after weaning (33). These cis-regulatory elements may be controlled by transcriptional enhancers or repressors. Indeed, we confirmed that 3 out of the 5 tested ACS that are located nearby genes important for proper *β*-cell function have a significant enhancer activity. In this study, we observed that accessible sites that were more open in adult islets seem to preexist in P10 islets, while sites less accessible in adults appear to be completely closed. Moreover, we found that preexisting programs were switched off at TSS while repressed or poised genes were fully activated by closing a distal repressor site. Intriguingly, Ackermann and others (3, 9) pointed out that many poised genes in *α* cells are a signature of functional *β*-cells and, consequently, could be of use for *α*-to-*β* cell reprogramming. This suggests plasticity and specialization of the different cell types along maturation and demonstrates rewiring of transcriptional programs at the chromatin level. For instance, reprogramming of acinar cells into insulin-producing cells has been done using adenoviral gene delivery of PDX1, NGN3 and MafA (4). A subsequent study characterized these changes using RNA-seq and ChIP-qPCR and revealed that loss of REST combined with PDX1 expression leads to activation of endocrine genes and correlates with epigenetic modifications of the local chromatin (34). Thus, islet plasticity is of interest for future treatment of diabetes through the production of mature surrogate insulin-producing cells (35). Using two different computational approaches and the Jaspar PWM database, we could find known and unforeseen transcriptional regulators potentially involved in the maturation process. We identified many DNA-binding proteins affecting chromatin accessibility, including MAF, FOX, FOS/JUN, NRF, E2F, CTCF, RFX, SREB, NKX6, REL, MEIS, and TEAD. Several of these transcription factors have been already implicated in pancreas development and postnatal islet maturation (36). For instance, E2F1 is an established cell survival and proliferation activator in *β*-cells (37) and we have recently shown that this transcription factor controls the expression of the long non-coding RNA H19 and is profoundly down-regulated during the postnatal period. Moreover, we obtained evidence suggesting that E2F1 and H19 may contribute to the decrease in *β*-cell mass in the maternal low protein diet offspring model (24). One of the identified transcriptional regulators, Scrt1, which is highly enriched in repressed sites in adults is of particular interest. Scrt1 was previously shown to be implicated in brain development (38, 39). Here, we confirmed that *Scrt1* mRNA levels are higher in rat adult islets, in both *α*- and *β*-cells. Interestingly, a significant up-regulation of *Scrt1* in adult islet cells was also reported in a recent study analyzing age-dependent gene expression and chromatin changes in human (40). Next, using a set of small interfering RNAs targeting *Scrt1* (siScrt1), we demonstrated that it is involved in *β*-cell proliferation. Finally, we identified Scrt1 targets using RNA-seq in FACS-sorted adult *β*-cells and confirmed that Scrt1 has a significant impact on transcripts related to proliferation. In addition, we observed that Scrt1 controls NFATc1/2 genes and consequently may control the calcineurin/NFAT pathway, known to be an important regulator of *β*-cell growth and function (41, 42). Our functional enrichment analysis showed that Scrt1 also influences the expression of genes related to oxygen sensing and autophagy. Accordingly, previous studies have demonstrated a role for the hypoxia-inducible factor HIF1a in normal *β*-cell function (43), and altered *β*-cell autophagy in human T2DM (44).

In this study, ATAC-seq and the mRNA microarray analyses have been performed in whole islets and not in FACS-sorted cells or in single cells. Several recent studies have demonstrated a large heterogeneity of gene expression at a single cell level using smFISH or single cell RNA-seq (45–48). Taking advantage of the heterogeneity of gene expression together with single cell ATAC-seq could shed more light on the transcriptional control of islet maturation and the different cell types involved.

Overall, we produced a high-resolution map of chromatin accessible sites in islets of 10-day-old pups and adult rats. These genome wide accessibility maps are an important resource to study cis-regulation of gene expression along maturation. Using these maps, we discovered a new important transcriptional repressor implicated in the maturation process, namely Scrt1, which controls *β*-cell proliferation and function. Manipulations of the level or the activity of this transcriptional regulator may favor the development of new future approaches aiming at generating surrogate insulin-producing cells. A global understanding of the molecular mechanisms and the transcription factors involved in functional maturation will be seminal for the design of *β*-cell-based replacement strategies for the treatment of diabetes (35).

## Methods and Materials

### Animals

Male (200-250g) and pregnant Sprague-Dawley rats were obtained from Janvier laboratories (Le Genest St-Isle, France). After birth, male and female pups were nursed until sacrifice at P10. All procedures were performed in accordance with the Guidelines for the care and use of laboratory animals from the National Institutes of Health and were approved by the Swiss Research Councils and Veterinary Office.

### Islet isolation, dispersion and sorting

Islets were isolated by collagenase digestion of the pancreas (49) followed by separation from digested exocrine tissue using an histopaque density gradient. After isolation, the islets were hand-picked and incubated for 2h in RPMI 1640 GlutaMAX medium (Invitrogen) containing 11 mM glucose and 2 mM L-glutamine and supplemented with 10% fetal calf serum (Gibco), 10 mM Hepes pH 7.4, 1 mM sodium pyruvate, 100 *µ*g/mL streptomycin and 100 IU/mL penicillin. Dissociated islet cells were obtained by incubating the islets in Ca^2+^/Mg^2+^ free phosphate buffered saline, 3 mM EGTA and 0.002% trypsin for 5 min at 37°C. For some experiments, islet cells were separated by Fluorescence-Activated Cell Sorting (FACS) based on *β*-cell autofluorescence, as previously described (50, 51). Sorted islet cells were seeded on plastic dishes coated with extracellular matrix secreted by 804 G rat bladder cancer cells (804 G ECM) (52). Enrichment of *α*- and *β*-cells was evaluated by double immunofluorescence staining using polyclonal guinea pig anti-insulin (dilution 1:40, PA1-26938 Invitrogen) and polyclonal mouse anti-glucagon (dilution 1:1000, Abcam Ab10988), followed by goat anti-guinea pig Alexa-Fluor-488 and goat anti-mouse Alexa-Fluor-555 (diluted 1:400, Thermofisher A11073 and A21422, respectively) secondary antibodies. On average, *β*- cell fractions contained 99.1 ± 0.9% insulin-positive cells and 0.6 ± 0.6% glucagon-positive cells and *α*-cell-enriched fractions contained 10.6 ± 8.2% insulin-positive cells and 88.8 ± 8.2% glucagon-positive cells.

### ATAC-seq sample preparation

ATAC-seq libraries were prepared as previously described (10) using dissociated islet cells from 3 adult male rats and from a mix of few P10 pups of 3 different litters. Briefly, 100’000 islet cells were re-suspended in 50 *µ*l of cold lysis buffer (10 mM Tris-HCl pH7.4, 10 mM NaCl, 3 mM MgCl_2_ and 0.1% IGEPAL CA-630) and centrifuged at 500g for 10 min at 4°C. The pellet was resuspended in the transposase reaction mix (Nextera kit, Illumina). The transposition reaction was performed at 37°C for 30 min and was followed by purification of the samples using the Qiagen MinElute PCR purification kit (Qiagen). Transposed DNA fragments were amplified for 11 cycles using the NEBnext high-fidelity PCR master mix and the Ad1_noMX and Ad2.1-2.6 barcoded primers from (10). Amplified libraries were purified with AMPure XP beads (Beckman Coulter) to remove contaminating primer dimers. Library quality was assessed using the Fragment Analyzer and quantitated using Qubit. All libraries were sequenced on Illumina HiSeq 2500 using 100 bp paired-end reads.

### ATAC-seq data quality control and analysis

Fastq files quality was assessed using FastQC (version 0.11.2) (53). Raw reads were aligned to the Rattus norvegicus reference genome assembly 5 (Rn5) using BWA (version 0.7.13) (54) with default settings. Quality control of the aligned reads was checked using Samstat (version 1.5) (55) and processed with Samtools (version 1.3) (56). Reads mapping to mitochondrial DNA were discarded from the analysis together with low quality reads (MAPQ < 30). Peak calling was performed in order to find accessible sites (ACSs) using Macs2 (version 2.1.1) (13) on adult and P10 samples concatenated separately. ACSs were then reunited in a single bed file and quantified using the pyDNase library (version 0.2.5) (57) (Table 1). We used FIMO (14) from the MEME suite (version 4.11.4) together with Jaspar 2016 position-weight matrices (19), to predict transcription factor binding sites.DeepTools (2.4.2) (58) was used to construct heatmaps around ACSs, TSS and TES. Finally, we used the R statistical software (version 3.4.2) and several bioconductor and CRAN packages to perform gene sets enrichment analysis (RDavidWebService, ClusterProfiler) (59), ACSs localization analysis (ChIPseeker) (16), motif enrichment analysis (FGSEA) (20) and penalized linear model analysis (GLMnet) (60) for motif selection. All sequencing tracks were viewed using the Integrated Genomic Viewer (IGV 2.4.8) (61). ATAC-seq raw data were deposited in GEO under the accession number GSE122747.

### mRNA Microarray

P10 and adult mRNA expression from (2, 24) were reanalyzed with EdgeR and RDavidWebservice (15, 62) (Figure 6-supplementary data table (6)). These microarray data are available in the GEO database under the accession number GSE106919.

### Cell line

The INS 832/13 rat *β*-cell line was provided by Dr. C. Newgard (Duke University) (63) and was cultured in RPMI 1640 GlutaMAX medium (Invitrogen) containing 11 mM glucose and 2 mM L-glutamine and supplemented with 10% fetal calf serum (Gibco), 10 mM Hepes pH 7.4, 1 mM sodium pyruvate and 0.05 mM of *β*-mercaptoethanol. INS 832/13 cells were cultured at 37°C in a humidified atmosphere (5% CO2, 95% air) and tested negative for mycoplasma contamination.

### Cell transfection

Dispersed rat islet cells or FACS-sorted *β*-cells were transfected with a pool of 4 siRNAs directed against rat Scrt1 or a negative control (On-Target plus 081299-02 and 001810-10 respectively, Dharmacon) using Lipofectamine RNAiMAX (Thermofisher). INS 832/13 cells were co-transfected with pGL3 promoter and psicheck plasmids (Promega) using lipofectamine 2000 (Thermofisher). pGL3 promoter vector was empty (control) or contained an enhancer region for *MafB, NeuroD1, Pax6* or *Syt4* (RNA synthesis, subcloning and plasmid sequencing were performed by GenScript, Netherlands) (Figure 3-Supplementary data (2)). RNA extraction and functional assays were performed 48h after transfection.

### Luciferase assay

Luciferase activities were measured in INS 832/13 cells using the Dual-Luciferase Reporter Assay System (Promega). Firefly luciferase activity was normalized to Renilla luciferase to minimize experimental variabilities. Experiments were performed in triplicates.

### RNA extraction, quantification and sequencing

RNA was extracted using miRNeasy micro kit (Qiagen) followed by DNase treatment (Promega). Gene expression levels were determined by qPCR using miScript II RT and SYBR Green PCR kits (Qiagen) and results were normalized to the housekeeping gene *Hprt1*. Data were analyzed using the 2^*−*ΔΔ*C*(*T*)^ method. Primer sequences are provided in Table 2.

**Table 2.**
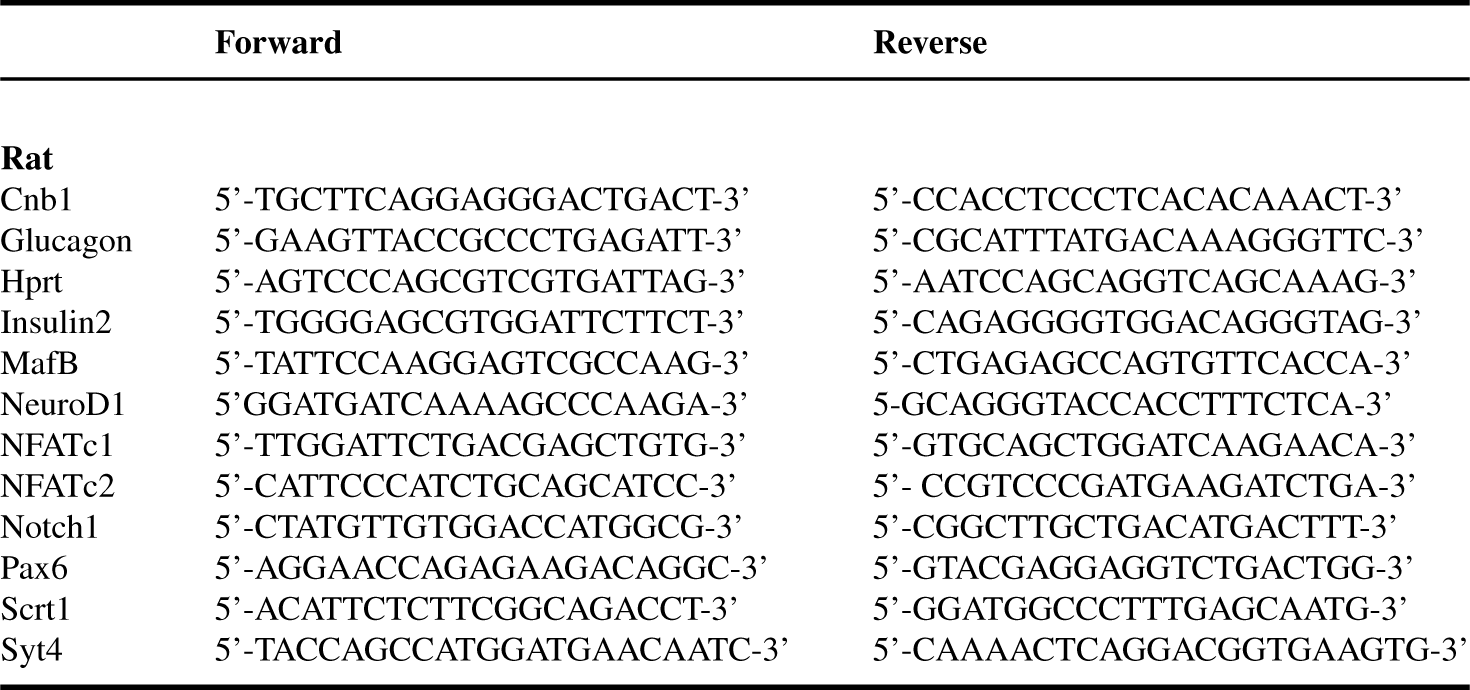
qPCR Primer Sequences

For mRNA-sequencing, the RNA was converted into a sequencing library using the Illumina TruSeq RNA-sequencing kit and standard Illumina protocols. Single-end, 151 nt long reads were obtained using a HiSeq 4000 instrument. Reads were aligned to the transcriptome (Rnor6) with Kallisto (64) and presented as transcripts per million (TPM) and EST pseudo counts (Additional data table (5)). Subsequently, differential expression was calculated using Sleuth (65). Finally, biological function and pathway analyses were performed using Cluster profiler (59). RNA-seq raw data were deposited in the GEO database under the accession number GSE130651.

### Insulin secretion and content

Transfected rat islet cells were pre-incubated for 30 min at 37°C in Krebs-Ringer bicarbonate buffer (KRBH) containing 2 mM glucose, 25 mM HEPES, pH 7.4 and 0.1 % BSA (Sigma-Aldrich), followed by 45 min incubation at 2 or 20 mM glucose. At the end of the incubation period, media were collected for insulin determination and rat islet cells were lysed with acid-ethanol (0.2 mM HCl in 75% ethanol) to extract total insulin content or with protein lysis buffer to measure total protein content (Bradford, BioRad). The amount of insulin released in the medium and remaining in the cells was measured by insulin Elisa kit (Mercodia). All experiments were performed in triplicates.

### Cell death assay

TUNEL staining on rat *β*-cells was performed 48h after transfection using the TMR red In Situ Cell Death Detection Kit (Roche) combined to polyclonal guinea pig anti-insulin (dilution 1:40, PA1-26938 Invitrogen) followed by incubation with goat anti-guinea-pig AlexaFluor 488 antibody (dilution 1:400, A11073 Thermofisher). Cell nuclei were stained with Hoechst 33342 (1 *µ*g/ml, Invitrogen). Coverslips were mounted on microscope glass slides with Fluor-Save mounting medium (VWR International SA) and were visualized with a Zeiss Axiovision fluorescence microscope. A minimum of 1 *\**10^3^ cells were counted per condition. Incubation for 24h with a mix pro-inflammatory cytokines (1 ng/mL IL-1*β*, 10 ng/mL TNF-*α* and 30 ng/mL IFN-*γ*) was used as positive control. Experiments were performed in single replicates.

### Proliferation assay

Transfected islet cells cultured on poly-L-lysine coated glass coverslips were fixed with ice-cold methanol and permeabilized with 0.5% (wt/vol) saponin (Sigma-Aldrich). The coverslips were first incubated with antibodies against Ki67 (dilution 1:500, Ab66155 Abcam) and polyclonal guinea pig anti-insulin (dilution 1:40, PA1-26938, Invitrogen) followed by incubation with anti-rabbit Alexa-Fluor-488 (dilution 1:400, A11008 Thermofisher) and anti-guinea-pig Alexa-Fluor-555 (dilution 1:400, A21435 Thermofisher) antibodies. At the end of the incubation, nuclei were stained with Hoechst 33342 (Invitrogen). Coverslips were mounted on microscope glass slides with Fluor-Save mounting medium (VWR International SA) and were visualized with a Zeiss Axiovision fluorescence microscope. Images of at least 1 *\** 10^3^ cells per condition were collected. Incubation with Prolactin (PRL 500 ng/ml during 48h) was used as positive control. Experiments were performed in single replicates.

### Statistical analysis

Data are expressed as mean ± SD. Statistical significance was determined using parametric unpaired two-tailed Student’s t-test or, for multiple comparisons, with one-way analysis of variance (ANOVA) of the means, followed by post-hoc Dunnett or Tukey test (Graph Pad Prism6). P-values less than 0.05 (p < 0.05) were considered statistically significant.

### Data availability

ATAC-seq raw data were deposited in GEO under accession number GSE122747. Microarray data from (2, 24) are available in the GEO database under accession number GSE106919. RNA-seq raw data were deposited in the GEO database under accession number GSE130651.

## Acknowledgments

We thank Charlotte Horr and Antonio Carlos Alves Meireles Filho for insightful discussions regarding the ATAC-seq sample preparation. This work was supported by the Swiss National Science Foundation (310030-169480 (RR)). We are grateful to the Placide Nicod fundation for their financial support (JS).

## SI datasets

1. ATAC-seq P10 Adult islets: Annotated ACS full table detected in pancreatic islet cells.
2. Validated ACS luciferase assay: Accessible sites information and sequence used in the luciferase assay to asses enhancer activity
3. TFBS ACS fgsea: FGSEA for motif accessibility. Table includes motif name, p-value, fdr, enrichment and normalized enrichment score, most extreme ACS rank, and number of target ACS significantly changing
4. Glmnet table: Penalized linear model for motif accessibility inference. Table include motif name, inferred accessibility coefficient *β*, motif consensus sequence, motif information content, and number of ACS targets
5. siSCRT1 vs Ctrl RNA-seq: SiScrt1 versus Ctrl RNA-seq of *β*-cells. The file contains the read counts per genes, and the statistical analysis of the differential gene expression performed with Sleuth.
6. Microarray p10 adult: Microarray of P10 versus adult rat pancreatic islet cells. The file contains the probe intensities and the statistical analysis of the differential gene expression performed with edgeR.
7. Sigdiff genes siSCRT1 Ctrl p10 adult: Genes differentially expressed in siScrt1 versus control adult rat *β*-cells (RNA-seq) and in adult versus P10 rat islets (microarray)
8. GOBPenrichment analysis siSCRT1 Ctrl p10 adult: Gene ontology of biological process enriched in differentially expressed genes in siScrt1 versus control adult rat *β*-cells and in adult versus P10 rat islets

---

**Fig. S1.**
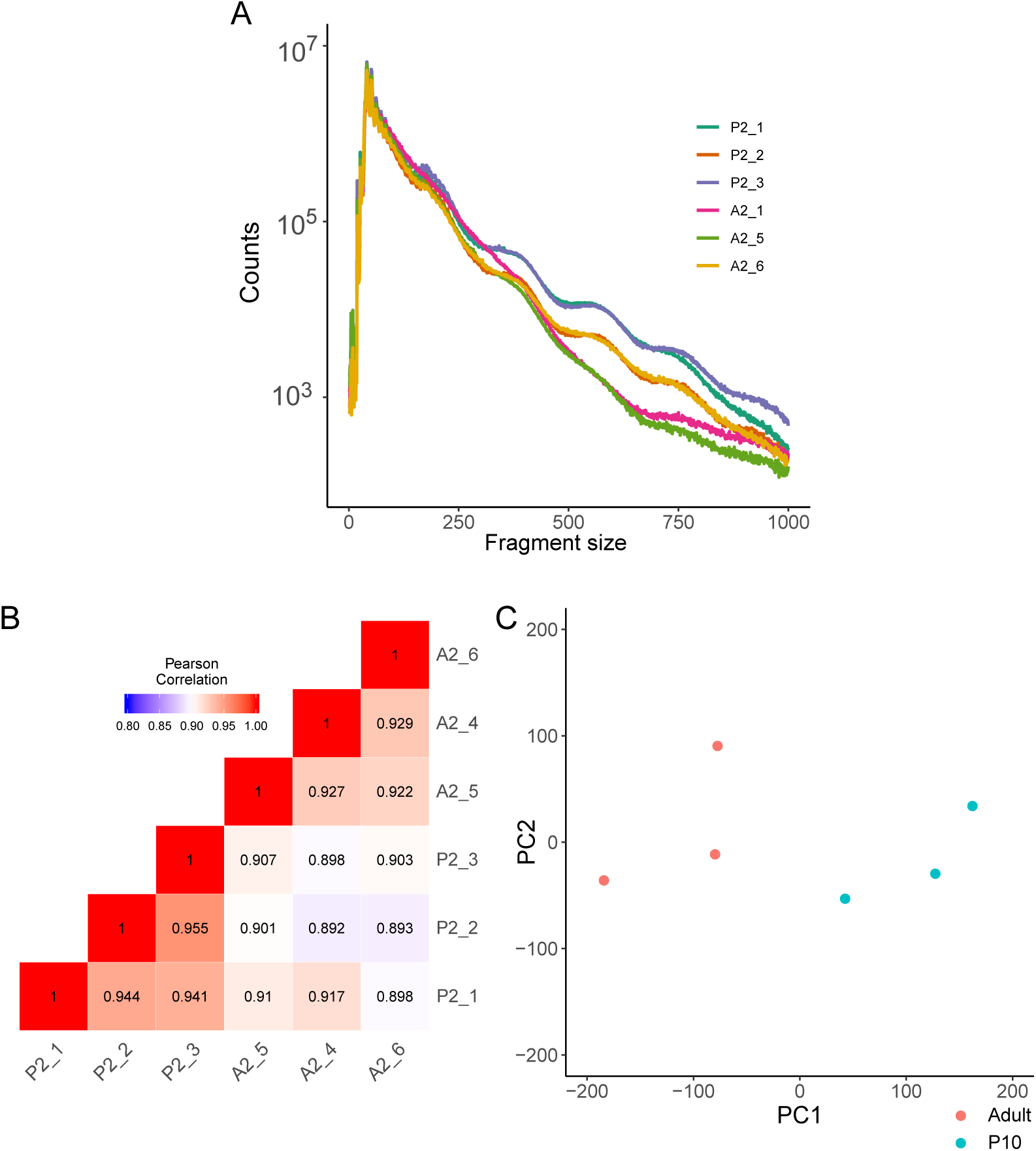
ATAC-seq quality control. (A) Fragment size distribution for each sample. (B) Correlation heatmap of samples. (C) PCA of each ATAC-seq sample.

**Fig. S2.**
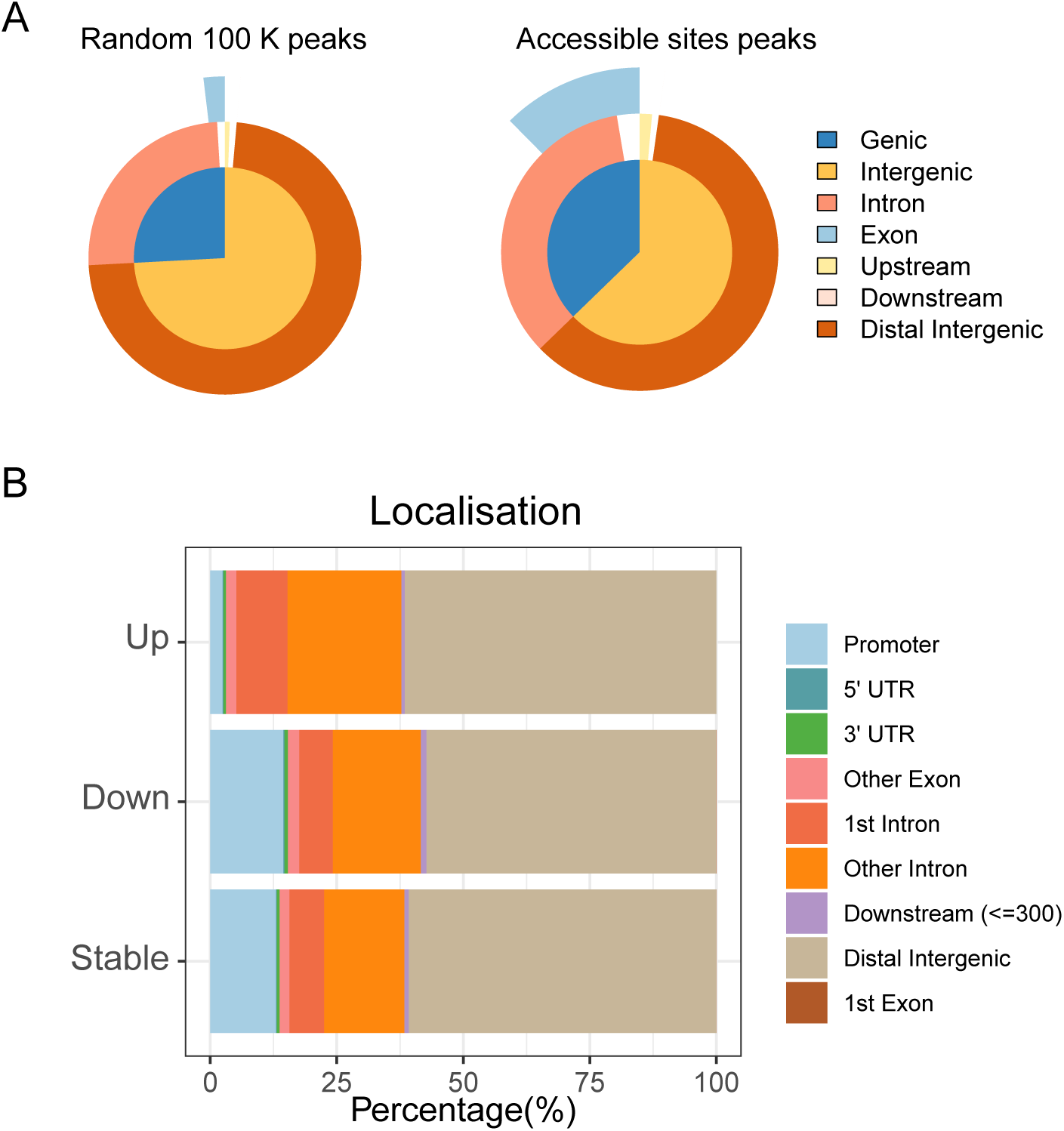
Genome-wide localisation and classification of ACS. (A) localisation of 100’000 random peaks compared to true accessible sites. True ACS are enriched in intronic and exonic regions. (B) localisation of stable ACS, more accessible ACS in adults (Up) and more accessible ACS in P10 (down). ACS more accessible in adults are enriched in introns, while stable and ACS more accessible in P10 are enriched in promoter regions.

**Fig. S3.**
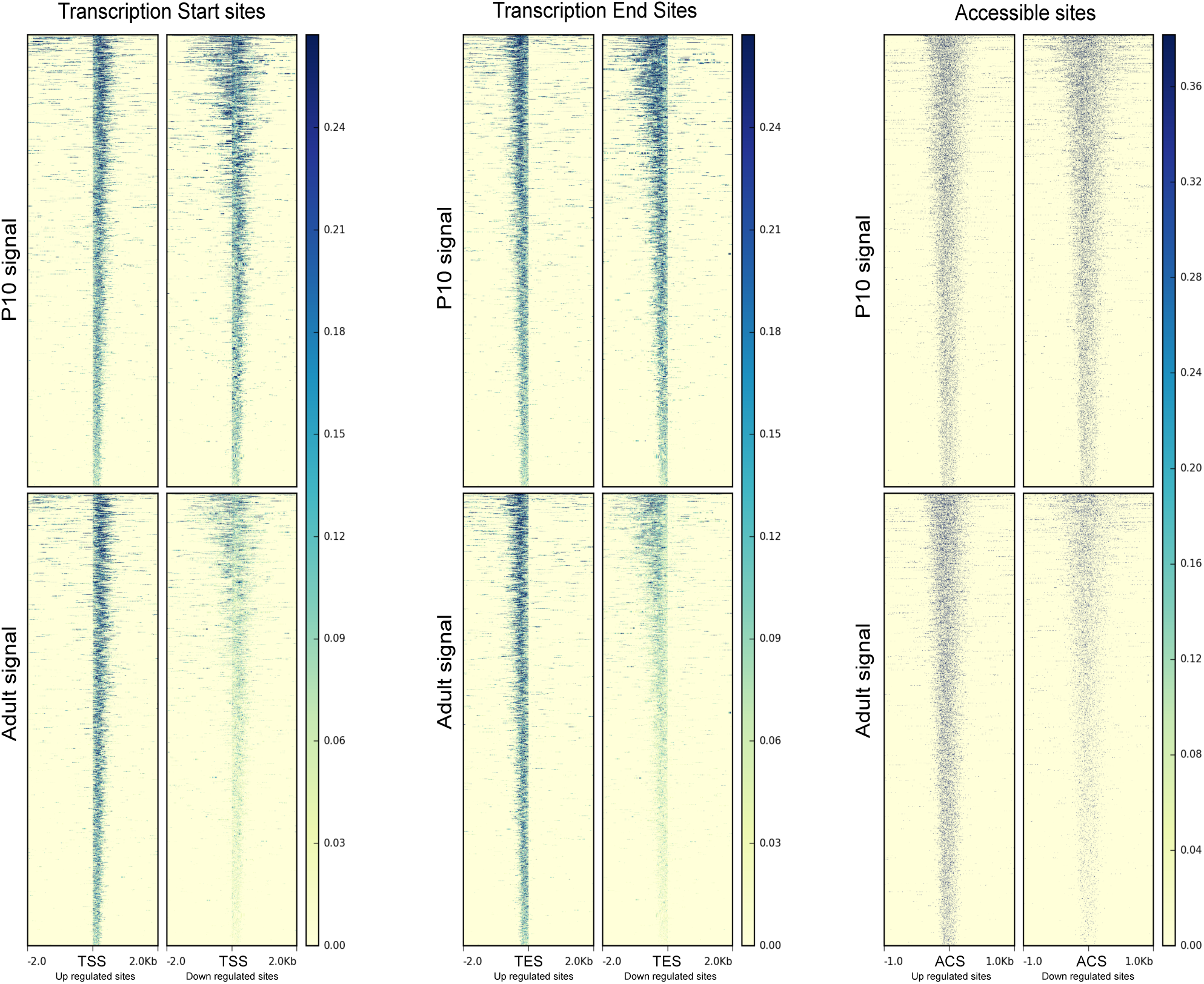
ATAC-seq signal is enriched at transcription start sites (TSS) and transcription end sites (TES) of genes nearby significantly changing ACS. The panel in the left represent TSS (+/- 2Kb) nearby up-regulated ACS or down-regulated ACS and in P10 or in adult condition (signal). Each line of the heatmap represent a single site. The middle panel show the same representation at TES. In the right panel, we show the signal around ACS center (+/- 1Kb). The ATAC-seq signal in the up-regulated ACS is slightly higher, while in the down-regulated ACS, we observed a much stronger difference between the two conditions.

**Fig. S4.**
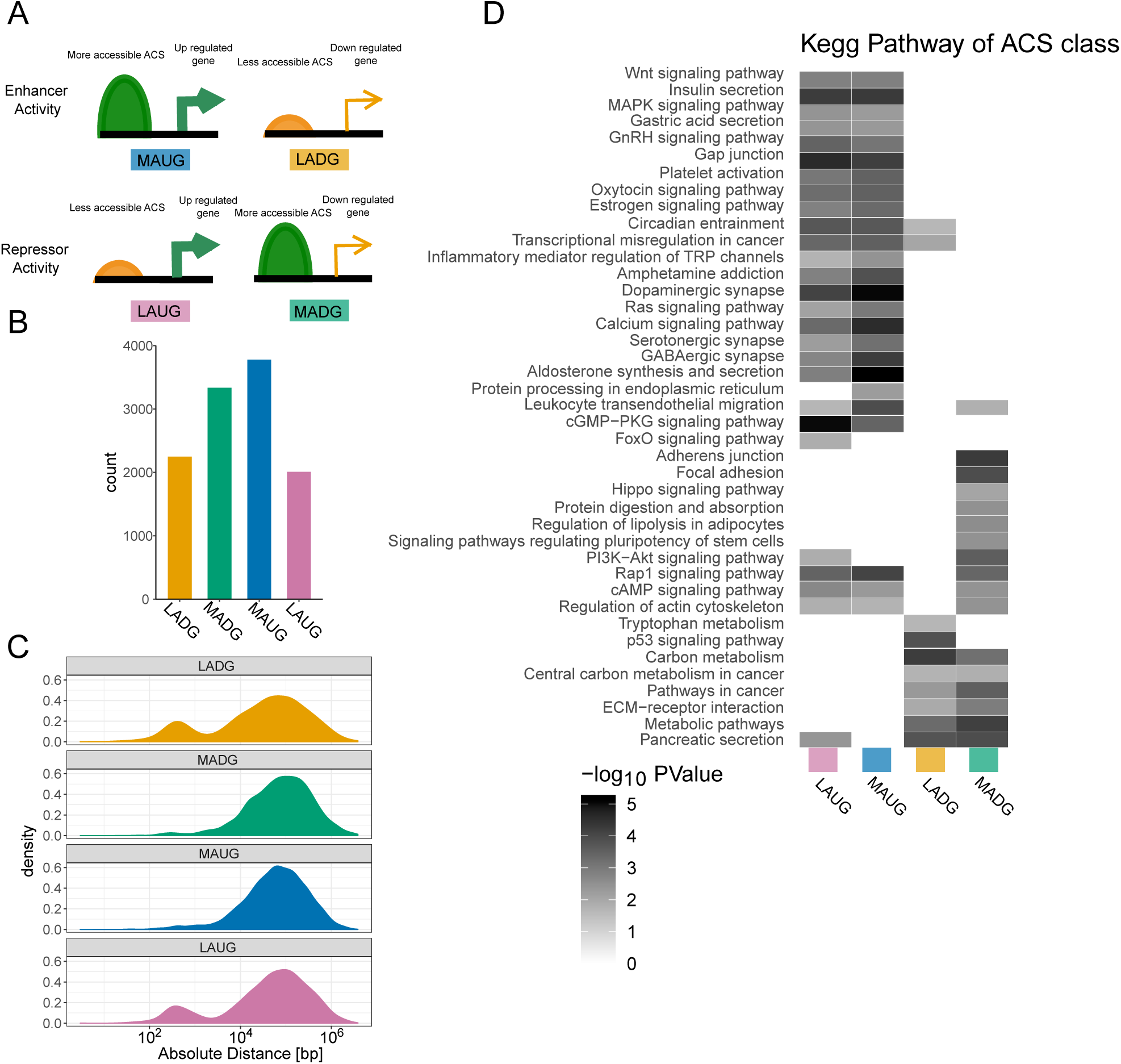
ACS localisation with respect to the closest gene, classification and pathway enrichment. (A) Scheme of ACS classification. (B) ACS count in each class and (C) absolute distance in log_10_ scale to the TSS of significantly differentially regulated mRNA between P10 and adult islets based on microarray for each class.(D) KEGG pathway analysis for each ACS class.

**Fig. S5.**
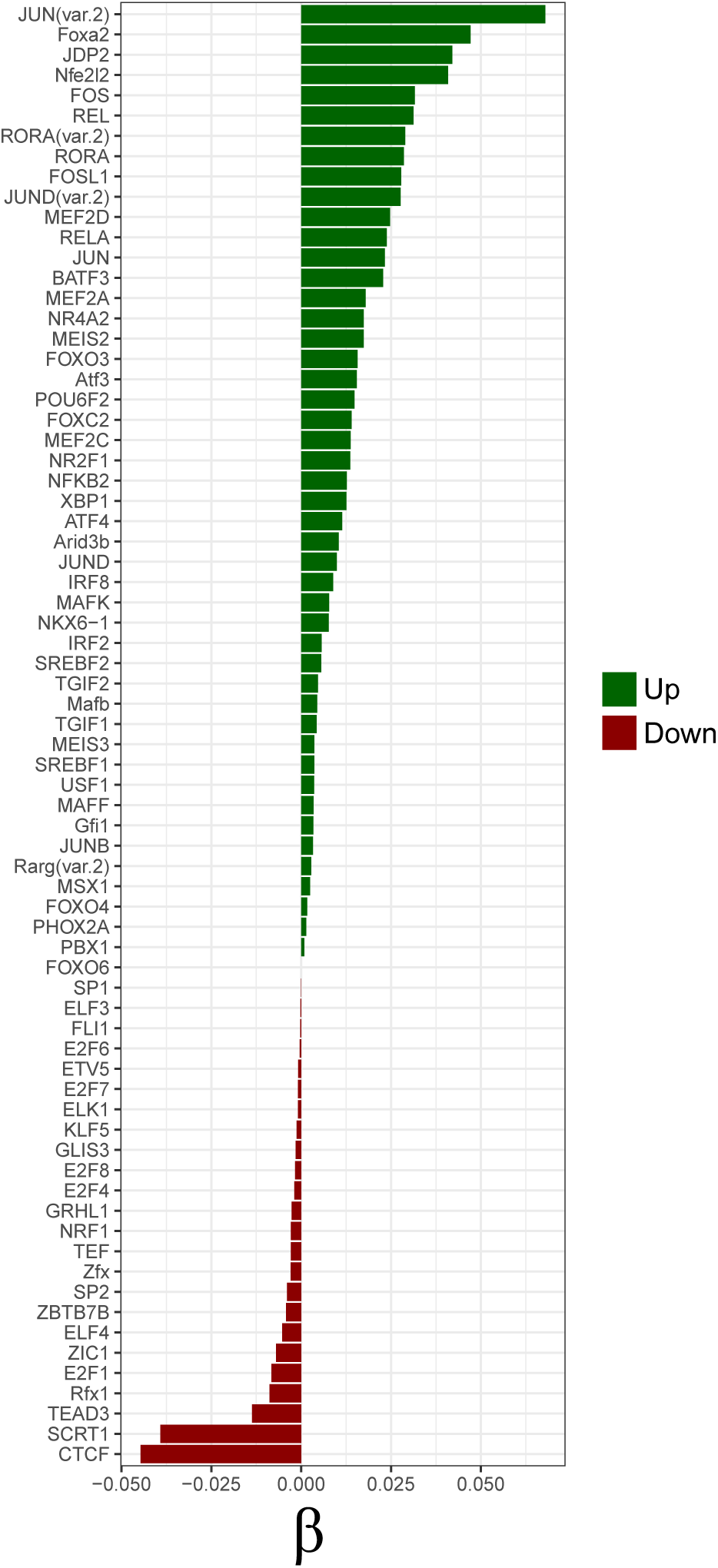
Penalised linear model GLMnet for transcription factor binding site motif activity inference. ACS sequences were scanned in order to construct the matrix of motifs per ACS. Every ACS with their respective log_2_ fold change (Adult/P10) were used to infer the motif activity, called *β* (see Methods). If *β* is negative, the motif is more accessible in P10 sites, while if the *β* is positive the motif explains a higher accessibility of the ACS in adults. Our linear model uses a penalty *λ* of 0.007 and an *α* of 0.1.

**Fig. S6.**
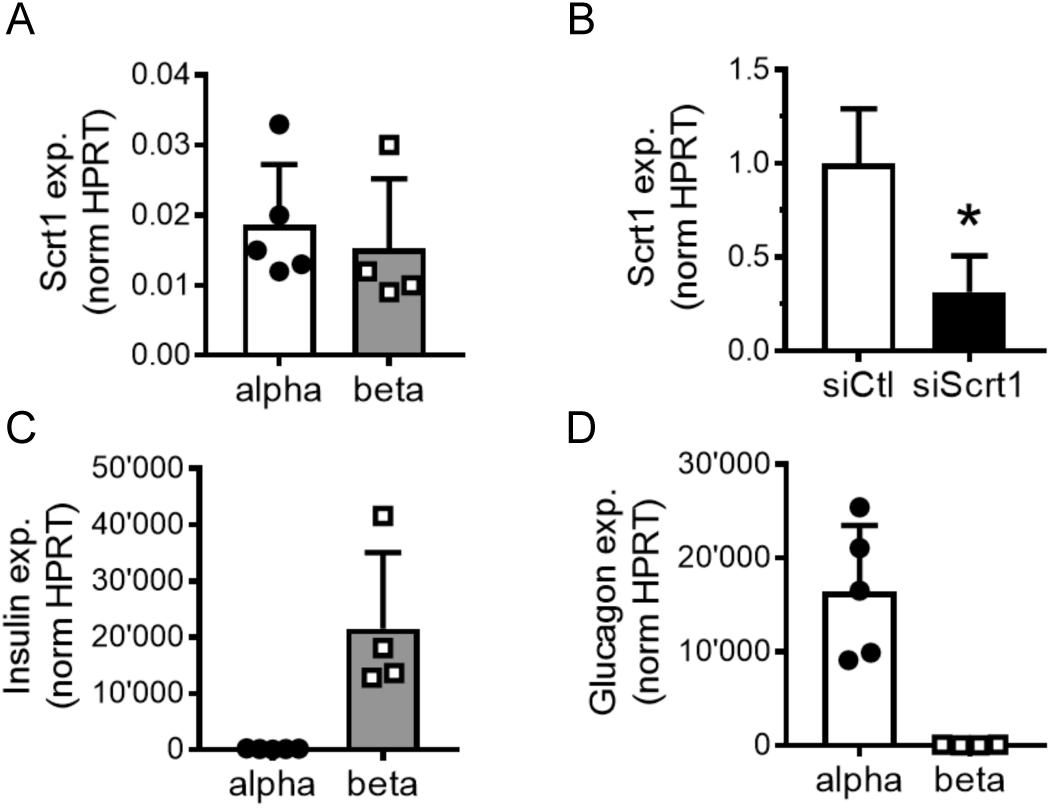
Downregulation of *Scrt1* expression in FACS-sorted *β*-cells. A-D) Indicated gene level was measured by qPCR in sorted *α*- and/or *β*-cells and normalized to *Hprt1* level. (A) *Scrt1* expression in sorted *α*- and *β*-cells. (B) *Scrt1* expression in FACS-sorted *β* cells 48h after transfection with a control siRNA (siCtl) or with siRNAs directed against Scrt1 (siScrt1). (C) Insulin and (D) glucagon expression in FACS-sorted *α*- and *β*-cell fractions. * p < 0.05 by Student’s t-test

**Fig. S7.**
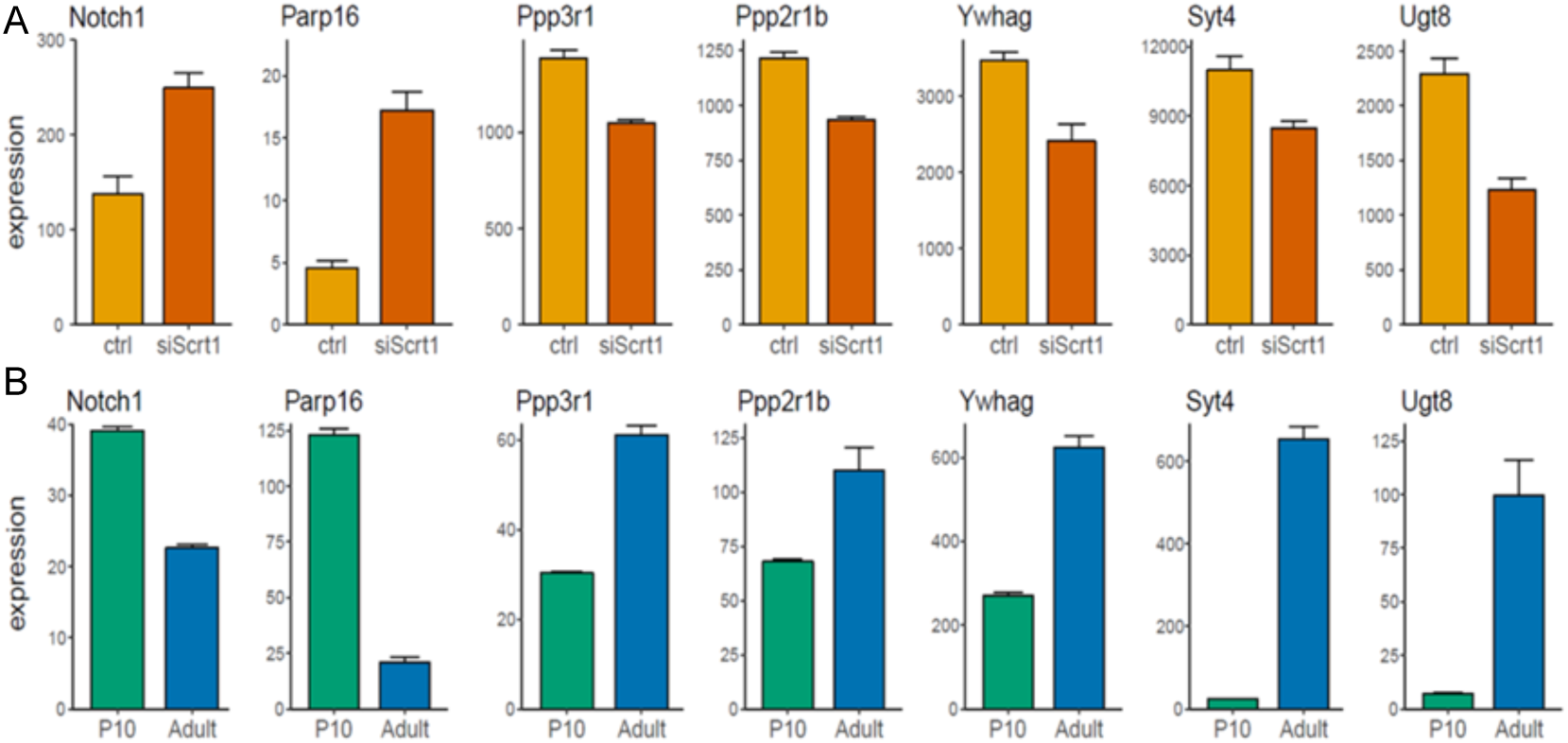
Expression changes in representative genes differentially expressed (FDR-adjusted p-value < 0.05) in (A) siScrt1 versus siRNA control (ctl) adult rat *β*-cells (RNA-seq) and in (B) P10 versus adult rat islets (micro-array).

## References

1. Miri Stolovich-Rain, Jonatan Enk, Jonas Vikesa, Finn Cilius Nielsen, Ann Saada, Benjamin Glaser, and Yuval Dor. Weaning triggers a maturation step of pancreatic *β* cells. Dev Cell, 32:535–545, 2015. ISSN 1534-5807. doi: 10.1016/j.devcel.2015.01.002.

2. Cécile Jacovetti, Scot J. Matkovich, Adriana Rodriguez-Trejo, Claudiane Guay, and Romano Regazzi. Postnatal *β*-cell maturation is associated with islet-specific microrna changes induced by nutrient shifts at weaning. Nat Commun, 6, 2015. ISSN 2041-1723. doi: 10.1038/ncomms9084.

3. Nuria C Bramswig, Logan J Everett, Jonathan Schug, Craig Dorrell, Chengyang Liu, Yanping Luo, Philip R Streeter, Ali Naji, Markus Grompe, and Klaus H Kaestner. Epigenomic plasticity enables human pancreatic *α* to *β* cell reprogramming. The Journal of clinical investigation, 123(3):1275–1284, 2013.

4. Luc Baeyens, Marie Lemper, Gunter Leuckx, Sofie De Groef, Paola Bonfanti, Geert Stangé, Ruth Shemer, Christoffer Nord, David W Scheel, Fong C Pan, et al. Transient cytokine treatment induces acinar cell reprogramming and regenerates functional beta cell mass in diabetic mice. Nature biotechnology, 32(1):76, 2014.

5. Carlos Larque, Myrian Velasco, Francisco Barajas-Olmos, Neyvis Garcia-Delgado, Juan Pablo Chávez-Maldonado, Jazmin Garcia-Morales, Lorena Orozco, and Marcia Hiriart. Transcriptome landmarks of the functional maturity of rat beta-cells, from lactation to adulthood. Journal of molecular endocrinology, 57(1):45–59, 2016.

6. Yoshio Fujitani. Transcriptional regulation of pancreas development and β-cell function [review]. Endocr J., 64:477–486, 2017. ISSN 0918-8959. doi: 10.1507/endocrj.ej17-0098.

7. Veronica Astro and Antonio Adamo. Epigenetic control of endocrine pancreas differentiation in vitro: current knowledge and future perspectives. Frontiers in cell and developmental biology, 6:141, 2018.

8. Dana Avrahami and Klaus H. Kaestner. Epigenetic regulation of pancreas development and function. Semin Cell Dev Biol, 23:693–700, 2012. ISSN 1084-9521. doi: 10.1016/j.semcdb.2012.06.002.

9. Amanda M. Ackermann, Zhiping Wang, Jonathan Schug, Ali Naji, and Klaus H. Kaestner. Integration of atac-seq and rna-seq identifies human alpha cell and beta cell signature genes. Mol Metab, 5:233–244, 2016. ISSN 2212-8778. doi: 10.1016/j.molmet.2016.01.002.

10. Jason D. Buenrostro, Paul G. Giresi, Lisa C. Zaba, Howard Y. Chang, and William J. Greenleaf. Transposition of native chromatin for fast and sensitive epigenomic profiling of open chromatin, dna-binding proteins and nucleosome position. Nat Methods, 10:1213–1218, 2013. ISSN 1548-7091. doi: 10.1038/nmeth.2688.

11. Jonathan Aryeh Sobel, Irina Krier, Teemu Andersin, Sunil Raghav, Donatella Canella, Federica Gilardi, Alexandra Styliani Kalantzi, Guillaume Rey, Benjamin Weger, Frederic Gachon, Matteo Dal Peraro, Nouria Hernandez, Ueli Schibler, Bart Deplancke, and Felix Naef and. Transcriptional regulatory logic of the diurnal cycle in the mouse liver. PLoS Biol., 2017. doi: 10.1101/077818.

12. Jason D Buenrostro, Beijing Wu, Howard Y Chang, and William J Greenleaf. Atac-seq: a method for assaying chromatin accessibility genome-wide. Current protocols in molecular biology, 109(1):21–29, 2015.

13. Yong Zhang, Tao Liu, Clifford A. Meyer, Jérôme Eeckhoute, David S. Johnson, Bradley E. Bernstein, Chad Nusbaum, Richard M. Myers, Myles Brown, Wei Li, and X. Shirley Liu. Model-based analysis of chip-seq (macs). Genome Biology, 9(9):R137, Sep 2008. ISSN 1474-760X. doi: 10.1186/gb-2008-9-9-r137.

14. Charles E. Grant, Timothy L. Bailey, and William Stafford Noble. FIMO: scanning for occurrences of a given motif. Bioinformatics, 27(7):1017–1018, 02 2011. ISSN 1367-4803. doi: 10.1093/bioinformatics/btr064.

15. Mark D Robinson, Davis J McCarthy, and Gordon K Smyth. edger: a bioconductor package for differential expression analysis of digital gene expression data. Bioinformatics, 26(1): 139–140, 2010.

16. Guangchuang Yu, Li-Gen Wang, and Qing-Yu He. ChIPseeker: an R/Bioconductor package for ChIP peak annotation, comparison and visualization. Bioinformatics, 31(14):2382–2383, 03 2015. ISSN 1367-4803. doi: 10.1093/bioinformatics/btv145.

17. Vanja Haberle and Alexander Stark. Eukaryotic core promoters and the functional basis of transcription initiation. Nature Reviews Molecular Cell Biology, 19(10):621, 2018.

18. Lorenzo Pasquali, Kyle J. Gaulton, Santiago A. Rodríguez-Seguí, Loris Mularoni, Irene Miguel-Escalada, İldem Akerman, Juan J. Tena, Ignasi Morán, Carlos Gómez-Marín, Martijn van de Bunt, Joan Ponsa-Cobas, Natalia Castro, Takao Nammo, Inês Cebola, Javier García-Hurtado, Miguel Angel Maestro, François Pattou, Lorenzo Piemonti, Thierry Berney, Anna L. Gloyn, Philippe Ravassard, José Luis Gómez Skarmeta, Ferenc Müller, Mark I. McCarthy, and Jorge Ferrer. Pancreatic islet enhancer clusters enriched in type 2 diabetes risk-associated variants. Nature Genetics, 46:136–143, 2014. ISSN 1061-4036. doi: 10.1038/ng.2870.

19. Albin Sandelin, Wynand Alkema, Pär Engström, Wyeth W Wasserman, and Boris Lenhard. Jaspar: an open-access database for eukaryotic transcription factor binding profiles. Nucleic acids research, 32(suppl_1):D91–D94, 2004.

20. Alexey Sergushichev. An algorithm for fast preranked gene set enrichment analysis using cumulative statistic calculation. BioRxiv, page 060012, 2016.

21. Jerome Friedman, Trevor Hastie, and Robert Tibshirani. Regularization paths for generalized linear models via coordinate descent. Journal of Statistical Software, 33(1):1–22, 2010.

22. Chris Lauber, Barbara Klink, and Michael Seifert. Comparative analysis of histologically classified oligodendrogliomas reveals characteristic molecular differences between sub-groups. BMC cancer, 18(1):399, 2018.

23. Taito Matsuda, Takashi Irie, Shutaro Katsurabayashi, Yoshinori Hayashi, Tatsuya Nagai, Nobuhiko Hamazaki, Aliya Mari D Adefuin, Fumihito Miura, Takashi Ito, Hiroshi Kimura, et al. Pioneer factor neurod1 rearranges transcriptional and epigenetic profiles to execute microglia-neuron conversion. Neuron, 101(3):472–485, 2019.

24. Clara Sanchez-Parra, Cécile Jacovetti, Olivier Dumortier, Kailun Lee, Marie-Line Peyot, Claudiane Guay, Marc Prentki, D Ross Laybutt, Emmanuel Van Obberghen, and Romano Regazzi. Contribution of the long noncoding rna h19 to *β*-cell mass expansion in neonatal and adult rodents. Diabetes, 67(11):2254–2267, 2018.

25. Kyle J Gaulton, Takao Nammo, Lorenzo Pasquali, Jeremy M Simon, Paul G Giresi, Marie P Fogarty, Tami M Panhuis, Piotr Mieczkowski, Antonio Secchi, Domenico Bosco, et al. A map of open chromatin in human pancreatic islets. Nature genetics, 42(3):255, 2010.

26. Arushi Varshney, Laura J Scott, Ryan P Welch, Michael R Erdos, Peter S Chines, Narisu Narisu, Ricardo D’O Albanus, Peter Orchard, Brooke N Wolford, Romy Kursawe, et al. Genetic regulatory signatures underlying islet gene expression and type 2 diabetes. Proceedings of the National Academy of Sciences, 114(9):2301–2306, 2017.

27. Matthias Thurner, Martijn Van De Bunt, Jason M Torres, Anubha Mahajan, Vibe Nylander, Amanda J Bennett, Kyle J Gaulton, Amy Barrett, Carla Burrows, Christopher G Bell, et al. Integration of human pancreatic islet genomic data refines regulatory mechanisms at type 2 diabetes susceptibility loci. Elife, 7:e31977, 2018.

28. William W Greenwald, Joshua Chiou, Jian Yan, Yunjiang Qiu, Ning Dai, Allen Wang, Naoki Nariai, Anthony Aylward, Jee Yun Han, Nikita Kadakia, et al. Pancreatic islet chromatin accessibility and conformation reveals distal enhancer networks of type 2 diabetes risk. Nature Communications, 10(1):2078, 2019.

29. Madhusudhan Bysani, Rasmus Agren, Cajsa Davegårdh, Petr Volkov, Tina Rönn, Per Unneberg, Karl Bacos, and Charlotte Ling. Atac-seq reveals alterations in open chromatin in pancreatic islets from subjects with type 2 diabetes. Scientific Reports, 9(1):7785, 2019.

30. Martin Piskacek, Ludmila Zotova, Gábor Zsurka, and Rudolf J Schweyen. Conditional knockdown of hmrs2 results in loss of mitochondrial mg+ uptake and cell death. Journal of cellular and molecular medicine, 13(4):693–700, 2009.

31. Kei Hang K Chan, Sara A Chacko, Yiqing Song, Michele Cho, Charles B Eaton, Wen-Chih H Wu, and Simin Liu. Genetic variations in magnesium-related ion channels may affect diabetes risk among african american and hispanic american women. The Journal of nutrition, 145(3):418–424, 2015.

32. Carsten Schmitz, Francina Deason, and Anne-Laure Perraud. Molecular components of vertebrate mg 2+-homeostasis regulation. Magnesium research, 20(1):6–18, 2007.

33. Cécile Jacovetti, Adriana Rodriguez-Trejo, Claudiane Guay, Jonathan Sobel, Sonia Gattesco, Volodymyr Petrenko, Camille Saini, Charna Dibner, and Romano Regazzi. Micrornas modulate core-clock gene expression in pancreatic islets during early postnatal life in rats. Diabetologia, 60(10):2011–2020, 2017.

34. Ofer Elhanani, Tomer Meir Salame, Jonathan Sobel, Dena Leshkowitz, Itay Vaknin, Dror Kolodkin-Gal, and Michael D. Walker. Rest inhibits direct reprogramming of pancreatic exocrine to endocrine cells by preventing pdx1-mediated activation of endocrine genes. In preparation, 2019.

35. AL Márquez-Aguirre, AA Canales-Aguirre, E Padilla-Camberos, H Esquivel-Solis, and NE Díaz-Martínez. Development of the endocrine pancreas and novel strategies for *β*-cell mass restoration and diabetes therapy. Brazilian Journal of Medical and Biological Research, 48(9):765–776, 2015.

36. Elizabeth Conrad, Roland Stein, and Chad S Hunter. Revealing transcription factors during human pancreatic *β* cell development. Trends in Endocrinology & Metabolism, 25(8):407–414, 2014.

37. Lluis Fajas, Jean-Sébastien Annicotte, Stéphanie Miard, David Sarruf, Mitsuhiro Watanabe, and Johan Auwerx. Impaired pancreatic growth, *β* cell mass, and *β* cell function in e2f1–/– mice. The Journal of clinical investigation, 113(9):1288–1295, 2004.

38. Faustino Marín and M Angela Nieto. The expression of scratch genes in the developing and adult brain. Developmental dynamics: an official publication of the American Association of Anatomists, 235(9):2586–2591, 2006.

39. Yasuhiro Itoh, Yasunobu Moriyama, Tsuyoshi Hasegawa, Takaho A Endo, Tetsuro Toyoda, and Yukiko Gotoh. Scratch regulates neuronal migration onset via an epithelial-mesenchymal transition–like mechanism. Nature neuroscience, 16(4):416, 2013.

40. H Efsun Arda, Lingyu Li, Jennifer Tsai, Eduardo A Torre, Yenny Rosli, Heshan Peiris, Robert C Spitale, Chunhua Dai, Xueying Gu, Kun Qu, et al. Age-dependent pancreatic gene regulation reveals mechanisms governing human *β* cell function. Cell metabolism, 23 (5):909–920, 2016.

41. Jeremy J Heit, Åsa A Apelqvist, Xueying Gu, Monte M Winslow, Joel R Neilson, Gerald R Crabtree, and Seung K Kim. Calcineurin/nfat signalling regulates pancreatic *β*-cell growth and function. Nature, 443(7109):345, 2006.

42. William R Goodyer, Xueying Gu, Yinghua Liu, Rita Bottino, Gerald R Crabtree, and Seung K Kim. Neonatal *β* cell development in mice and humans is regulated by calcineurin/nfat. Developmental cell, 23(1):21–34, 2012.

43. Kim Cheng, Kenneth Ho, Rebecca Stokes, Christopher Scott, Sue Mei Lau, Wayne J Hawthorne, Philip J O’connell, Thomas Loudovaris, Thomas W Kay, Rohit N Kulkarni, et al. Hypoxia-inducible factor-1*α* regulates *β* cell function in mouse and human islets. The Journal of clinical investigation, 120(6):2171–2183, 2010.

44. M Masini, M Bugliani, R Lupi, S Del Guerra, Ugo Boggi, Franco Filipponi, Lorella Marselli, Pellegrino Masiello, and Piero Marchetti. Autophagy in human type 2 diabetes pancreatic beta cells. Diabetologia, 52(6):1083–1086, 2009.

45. Lydia Farack, Matan Golan, Adi Egozi, Nili Dezorella, Keren Bahar Halpern, Shani Ben-Moshe, Immacolata Garzilli, Beáta Tóth, Lior Roitman, Valery Krizhanovsky, et al. Transcriptional heterogeneity of beta cells in the intact pancreas. Developmental cell, 48(1): 115–125, 2019.

46. Wei-Lin Qiu, Yu-Wei Zhang, Ye Feng, Lin-Chen Li, Liu Yang, and Cheng-Ran Xu. Deciphering pancreatic islet *β* cell and *α* cell maturation pathways and characteristic features at the single-cell level. Cell metabolism, 25(5):1194–1205, 2017.

47. Nicole AJ Krentz, Michelle YY Lee, Eric E Xu, Shannon LJ Sproul, Alexandra Maslova, Shugo Sasaki, and Francis C Lynn. Single-cell transcriptome profiling of mouse and hesc-derived pancreatic progenitors. Stem cell reports, 11(6):1551–1564, 2018.

48. Chun Zeng, Francesca Mulas, Yinghui Sui, Tiffany Guan, Nathanael Miller, Yuliang Tan, Fenfen Liu, Wen Jin, Andrea C Carrano, Mark O Huising, et al. Pseudotemporal ordering of single cells reveals metabolic control of postnatal *β* cell proliferation. Cell metabolism, 25 (5):1160–1175, 2017.

49. Mitsukazu Gotoh, TAKASHI Maki, SUSUMU Satomi, JANIS Porter, SUSAN Bonner-Weir, Carl J O’hara, and Anthony P Monaco. Reproducible high yield of rat islets by stationary in vitro digestion following pancreatic ductal or portal venous collagenase injection. Transplantation, 43(5):725–730, 1987.

50. Claudiane Guay, Janine K. Kruit, Sophie Rome, Véronique Menoud, Niels L. Mulder, Angelika Jurdzinski, Francesca Mancarella, Guido Sebastiani, Alena Donda, Bryan J. Gonzalez, Camilla Jandus, Karim Bouzakri, Michel Pinget, Christian Boitard, Pedro Romero, Francesco Dotta, and Romano Regazzi. Lymphocyte-derived exosomal micrornas promote pancreatic *β* cell death and may contribute to type 1 diabetes development. Cell Metabolism, 29:348–361.e6, 2019. ISSN 1550-4131. doi: 10.1016/j.cmet.2018.09.011.

51. Martin Köhler, Elisabetta Daré, Muhammed Yusuf Ali, Subu Surendran Rajasekaran, Tilo Moede, Barbara Leibiger, Ingo B. Leibiger, Annika Tibell, Lisa Juntti-Berggren, and Per-Olof Berggren. One-step purification of functional human and rat pancreatic alpha cells. Integr Biol (Camb), 4:209, 2012. ISSN 1757-9694. doi: 10.1039/c2ib00125j.

52. G. Parnaud, D. Bosco, T. Berney, F. Pattou, J. Kerr-Conte, M. Y. Donath, C. Bruun, T. Mandrup-Poulsen, N. Billestrup, and P. A. Halban. Proliferation of sorted human and rat beta cells. Diabetologia, 51:91–100, 2007. ISSN 0012-186X. doi: 10.1007/s00125-007-0855-1.

53. S Andrews. Fastqc: a quality control tool for high throughput sequence data [internet]. Cambridge, UK: Babraham Bioinformatics, The Babraham Institute, 2010.

54. Heng Li and Richard Durbin. Fast and accurate short read alignment with burrows–wheeler transform. bioinformatics, 25(14):1754–1760, 2009.

55. Timo Lassmann, Yoshihide Hayashizaki, and Carsten O Daub. Samstat: monitoring biases in next generation sequencing data. Bioinformatics, 27(1):130–131, 2010.

56. Heng Li, Bob Handsaker, Alec Wysoker, Tim Fennell, Jue Ruan, Nils Homer, Gabor Marth, Goncalo Abecasis, and Richard Durbin. The sequence alignment/map format and samtools. Bioinformatics, 25(16):2078–2079, 2009.

57. Jason Piper, Markus C. Elze, Pierre Cauchy, Peter N. Cockerill, Constanze Bonifer, and Sascha Ott. Wellington: a novel method for the accurate identification of digital genomic footprints from dnase-seq data. Nucleic Acids Res, 41:e201–e201, 2013. ISSN 1362-4962. doi: 10.1093/nar/gkt850.

58. Fidel Ramírez, Friederike Dündar, Sarah Diehl, Björn A Grüning, and Thomas Manke. deep-tools: a 59. fl60. exible platform for exploring deep-sequencing data. Nucleic acids research, 42 (W1):W187–W191, 2014.

59. Guangchuang Yu, Li-Gen Wang, Yanyan Han, and Qing-Yu He. clusterprofiler: an r package for comparing biological themes among gene clusters. Omics: a journal of integrative biology, 16(5):284–287, May 2012. ISSN 1536-2310.

60. Jerome Friedman, Trevor Hastie, and Robert Tibshirani. Regularization paths for generalized linear models via coordinate descent. Journal of Statistical Software, 33, 2010. ISSN 1548-7660. doi: 10.18637/jss.v033.i01.

61. James T. Robinson, Helga Thorvaldsdóttir, Wendy Winckler, Mitchell Guttman, Eric S. Lander, Gad Getz, and Jill P. Mesirov. Integrative genomics viewer. Nature biotechnology, 29 (1):24–26, January 2011. ISSN 1087-0156.

62. C. Fresno and E. A. Fernandez. Rdavidwebservice: a versatile r interface to david. Bioinformatics, 29:2810–2811, 2013. ISSN 1367-4803. doi: 10.1093/bioinformatics/btt487.

63. Hans E Hohmeier, Hindrik Mulder, Guoxun Chen, Rosemarie Henkel-Rieger, Marc Prentki, and Christopher B Newgard. Isolation of ins-1-derived cell lines with robust atp-sensitive k+ channel-dependent and-independent glucose-stimulated insulin secretion. Diabetes, 49(3): 424–430, 2000.

64. Nicolas L Bray, Harold Pimentel, Páll Melsted, and Lior Pachter. Near-optimal probabilistic rna-seq quantification. Nature biotechnology, 34(5):525, 2016.

65. Harold Pimentel, Nicolas L Bray, Suzette Puente, Páll Melsted, and Lior Pachter. Differential analysis of rna-seq incorporating quantification uncertainty. Nature methods, 14(7):687, 2017.

